# HELLS and PRDM9 form a Pioneer Complex to Open Chromatin at Meiotic Recombination Hotspots

**DOI:** 10.1101/764183

**Authors:** Catrina Spruce, Sibongakonke Dlamini, Guruprasad Ananda, Naomi Bronkema, Hui Tian, Ken Paigen, Gregory W. Carter, Christopher L Baker

**Affiliations:** The Jackson Laboratory, Bar Harbor, ME 04660

**Keywords:** pioneer factor, chromatin remodeling, histone modification, nucleosome-depleted region, germ cells

## Abstract

Chromatin barriers prevent spurious interactions between regulatory elements and DNA-binding proteins. One such barrier, whose mechanism for overcoming is poorly understood, is access to recombination hotspots during meiosis. Here we show that the chromatin remodeler HELLS and DNA-binding protein PRDM9 function together to open chromatin at hotspots and provide access for the DNA double-strand break (DSB) machinery. Recombination hotspots are decorated by a unique combination of histone modifications, not found at other regulatory elements. HELLS is recruited to hotspots by PRDM9, and is necessary for both histone modifications and DNA accessibility at hotspots. In male mice lacking HELLS, DSBs are retargeted to other sites of open chromatin, leading to germ cell death and sterility. Together, these data provide a model for hotspot activation where HELLS and PRDM9 function as a pioneer complex to create a unique epigenomic environment of open chromatin, permitting correct placement and repair of DSBs.

## INTRODUCTION

In eukaryotic cells, DNA is normally wrapped around an octamer of histone proteins to form nucleosomes, the basic unit of chromatin. Chromatin functions as a physical barrier to DNA, and regulating access to DNA sequences in chromatin is an essential aspect to normal cellular function. Numerous proteins have evolved to carry out this function; these include pioneer factors that determine the location of open chromatin, chromatin remodelers that physically move or evict nucleosomes from DNA, and epigenetic writers that catalyze post-translational modifications on histones (Clapier and Cairns, 2009; Clapier et al., 2017; Iwafuchi-Doi and Zaret, 2014).

Meiosis, which is essential for all sexual reproduction, involves major reorganization of chromatin compartments (Alavattam et al., 2019; Patel et al., 2019), along with chromosome condensation, pairing, and ultimately genetic recombination. The outcome of recombination is the exchange of genetic material between parental chromosomes resulting in the shuffling of alleles between generations. In mammals, fungi, and plants, recombination is concentrated at specialized sites termed hotspots (Baudat et al., 2013; Paigen and Petkov, 2010; Tock and Henderson, 2018). The physical exchange of DNA is initiated when the topoisomerase SPO11 is recruited to hotspots, where it induces the double-strand breaks (DSBs) required for DNA exchange between homologous chromosomes (Keeney, 2001). Significant chromatin remodeling is a prerequisite for the proper targeting and access of SPO11 to hotspot DNA; this remodeling likely involves overcoming chromatin barriers that regulate DNA accessibility.

In most mammals, hotspot locations are determined by the DNA-binding protein PRDM9 (Baudat et al., 2010; Myers et al., 2010; Parvanov et al., 2010). PRDM9 contains a SET domain, that catalyzes both histone H3 lysine 4 trimethylation (H3K4me3) and H3K36me3 (Buard et al., 2009; Eram et al., 2014; Powers et al., 2016; Wu et al., 2013), and a series of tandem C-terminal zinc-finger domains that bind DNA. *Prdm9* is highly polymorphic with the majority of differences in regions encoding the zinc-finger domains (Berg et al., 2011; Kono et al., 2014; Schwartz et al., 2014). Allelic variants bind different DNA sequences, ultimately dictating the location of hotspots. PRDM9 binding occurs within nucleosome-depleted regions (NDR) (Baker et al., 2014; Baker et al., 2015a) that are subsequently targeted for DSBs by SPO11 (Baker et al., 2014; Lange et al., 2016). To understand how hotspots can occur in regions of otherwise closed chromatin, we postulated that PRDM9 functions as a pioneer factor to overcome chromatin barriers at hotspots.

Pioneer factors initiate formation of accessible chromatin by engaging partial DNA recognition motifs present within nucleosomal DNA. (Soufi et al., 2012; Zaret and Mango, 2016). Binding of pioneer leads to increased accessibility and depletion of nucleosomes, ultimately enabling other protein factors to gain access to regulatory elements (Iwafuchi-Doi and Zaret, 2014). Increased DNA accessibility is also accompanied by an increase in active histone modifications. Because of these unique properties, the study of pioneer factors have focused on their role as master regulators of cellular differentiation and lineage reprogramming (Soufi et al., 2012; Zaret and Mango, 2016).

Recruitment of chromatin-remodeling enzymes by pioneer factors is a critical step in creating accessible DNA (King and Klose, 2017; Swinstead et al., 2016b). The *Snf2* family of chromatin-remodelers utilize energy from ATP hydrolysis to exchange, evict, and reposition nucleosomes to create open chromatin (Clapier and Cairns, 2009; Flaus et al., 2006). Several chromatin-remodeling factors are essential for normal germ-cell development. Meiosis-specific loss of *Brg1*, *Hells* (*Lsh*), and *Ino80* each result in the persistence of unrepaired DSBs, incomplete synapsis of homologous chromosomes, and pachytene arrest in meiotic prophase I (Kim et al., 2012; Serber et al., 2016; Zeng et al., 2011). These phenotypes are similar to loss of *Prdm9*, indicating that these chromatin remodelers might play direct roles in nucleosome remodeling at recombination hotspots.

Given the broad similarity in requirements for accessible DNA for both pioneer factor-mediated control of gene regulation and meiotic recombination, we determined the mechanisms supporting PRDM9-mediated chromatin organization and remodeling at hotspots.

## RESULTS

### Hotspots are marked by a unique epigenomic state

To functionally annotate the chromatin landscape of male germ cells, we determined co-occupancy of histone modifications using ChromHMM (Ernst and Kellis, 2012). We incorporated new and previously published nucleosome-resolution ChIP-seq data from 12-14 day post-partum (dpp) C57BL/6J (B6) mice to generate an 11-state model (**Fig. 1A**, **Table S1**). This germ cell model recapitulated many of the previously observed chromatin states found in other cell types including promoters and insulators (states 1), enhancers (states 2-5) transcriptionally active regions (state 6), Polycomb repressive complex-repressed (state 8), and heterochromatin (states 9 and 10) (Fig. 1A and B).

**Figure 1.**
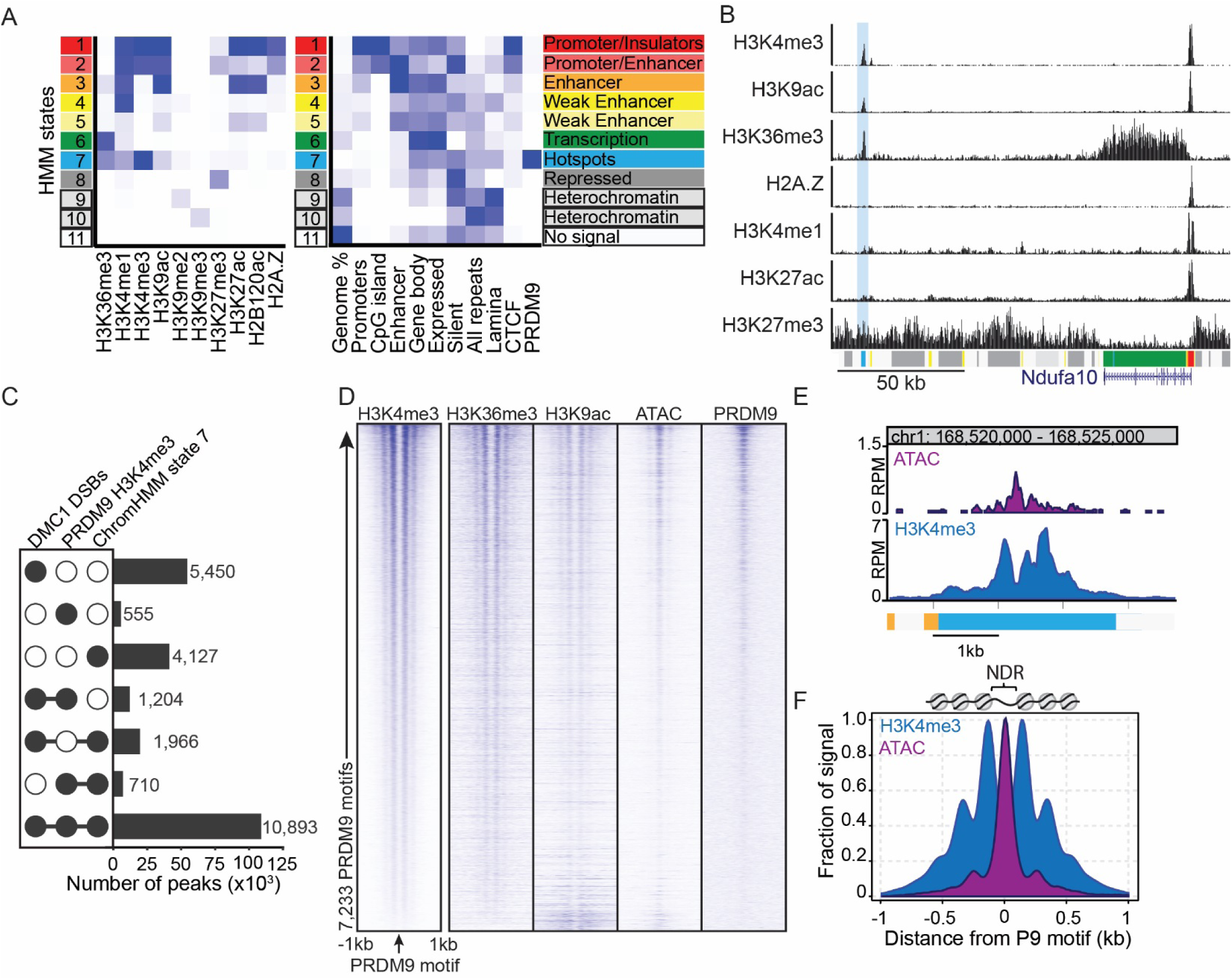
Recombination hotspots have a unique epigenomic signature and are sites of open chromatin. (A) *Left* – heat map of the emission parameters for 11-state ChromHMM solution based on ChIP-seq for 10 histone modifications. *Right* – heat map showing fold enrichment for indicated annotations for each state. (B) Example ChIP-seq profiles for data used to build ChromHMM model. *Bottom* – HMM genome annotations (blue highlight – example hotspot). (C) Upset plot showing overlap between meiotic DSBs (DMC1 ChIP), PRDM9-dependent H3K4me3, and HMM state 7. (D) Heat maps of histone modifications, ATAC-seq, and PRDM9 ChIP-seq spanning ±1kb of PRDM9 motif. Heat maps are normalized counts per million (cpm) in 10 bp bins. (E) Genomic profile for example locus at Pbx1 hotspot showing ATAC-seq signal. (F) Meta-profile of histone modifications and open chromatin at recombination hotspots from *D* anchored by PRDM9 motif background subtracted and scaled to fraction of the maximum value.

Of these eleven chromatin states, only state 7 was enriched for PRDM9 binding sites; it is distinguished by a unique combination of histone modifications. In addition to H3K4me3 and H3K36me3, state 7 is characterized by H3K4me1 and H3K9ac (Fig. 1A and B). In contrast to other acetylation marks often found at enhancers (i.e. H3K27ac and H2BK120ac), only H3K9ac is enriched at hotspots. The majority (78%, 14,773/18,900) of all state 7 locations overlap with previously reported locations of B6 meiosis-specific DSBs (Smagulova et al., 2016) and PRDM9-dependent H3K4me3 modification (Baker et al., 2014) (**Fig. 1C**), highlighting that state 7 represents recombination hotspots. Interestingly, compared to other phyla (Choi et al., 2013; Yamada et al., 2018), we did not detect H2A.Z at hotspots in mice. Together, these data identify a unique combination of histone features that biochemically distinguish recombination hotspots from other regulatory elements, and may be involved in creating chromatin accessibility for DSB formation.

### Hotspots are sites of accessible chromatin

Given that recombination hotspots are marked by histone modifications associated with active chromatin, we measured DNA accessibility in germ cells using the assay for transposase-accessible chromatin (ATAC-seq) (Buenrostro et al., 2013). Biological replicates proved highly reproducible (r = 0.96), showed enrichment of open chromatin at promoters, and revealed typical nucleosome banding patterns (**Fig. S1A-C)**, all indicators of high-quality libraries. At PRDM9-dependent H3K4me3 sites, we detected increased DNA accessibility overlapping with PRDM9 motif locations (Fig. 1D-F). Overall, ATAC-seq identified fewer hotspots compared to H3K4me3 ChIP-seq (2,902 ATAC peaks overlapping 13,498 PRDM9-dependent H3K4me3).

Hotspots with greater PRDM9 binding and histone modification generally have more open chromatin (**Fig. 1D**). On average, DNA accessibility showed an inverse relationship to nucleosome positions at hotspots (Fig. 1E and F). Filtering ATAC-seq reads for those that fall within NDRs (i.e. < 120 bp) (Buenrostro et al., 2013), showed that open chromatin was highest where PRDM9 binds, although the adjacent inter-nucleosome regions also showed increased accessibility. Together, these data build on an earlier observation that detected increased DNase hypersensitivity at a single hotspot (Shenkar et al., 1991), and demonstrate that generally hotspots are sites of open chromatin.

### PRDM9 acts as a meiosis-specific pioneer factor to create open chromatin

We next applied three genetic tests to determine whether the unique epigenomic state and presence of open chromatin at hotspots is PRDM9-dependent: 1) determination of chromatin accessibility in spermatocytes with different *Prdm9* alleles but controlled overall genetic background, 2) assessment of chromatin with the same *Prdm9* allele and but different genetic backgrounds, and 3) determination of chromatin states in heterozygous F1 hybrids with different *Prdm9* alleles and different genetic backgrounds.

To maintain the same genetic background with different *Prdm9* alleles, we used a ‘knock-in’ (KI) mouse strain that exchanges the endogenous B6 *Prdm9* zinc-finger domain with the one from CAST/EiJ mice (Cst) (Baker et al., 2014). We measured H3K9ac and chromatin accessibility in enriched KI germ cells (**Fig. S1D**), combined with H3K4me3 (Baker et al., 2014), and compared these to the B6 results (Fig. 2A and B**, Fig. S1D-H**). Quantifying differences between B6 and KI, we identified 5,775 ATAC peaks and 14,249 H3K9ac peaks increased in KI (FDR < 0.01). Of these, 92% of the ATAC peaks (n = 5,305) and 94% of the H3K9ac peaks (n = 13,408) overlap with reported *Prdm9^Cst^-*dependent H3K4me3 peaks (Baker et al., 2014).

**Figure 2.**
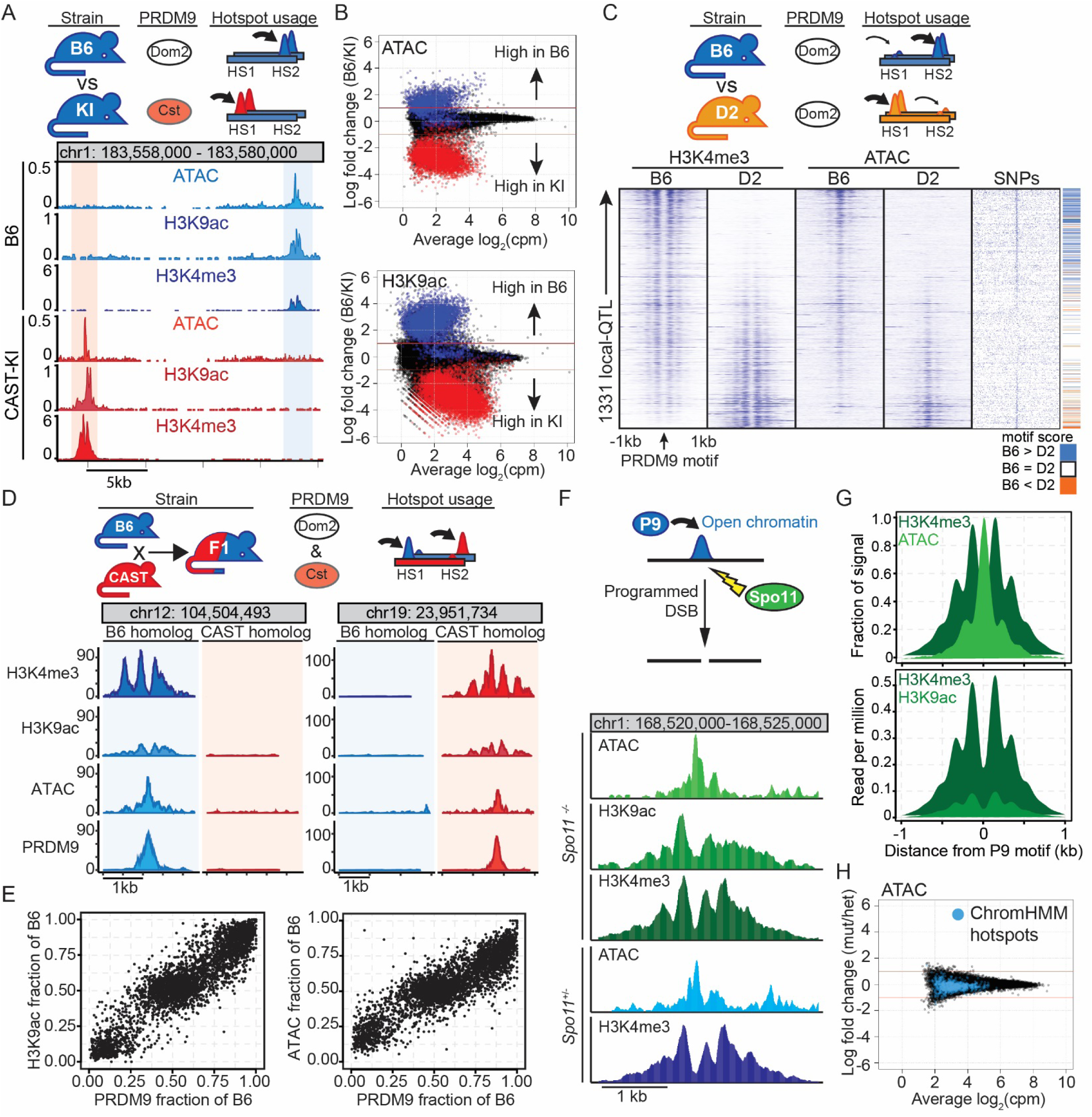
Open chromatin at hotspots is dependent on PRDM9 but not on DSBs. (A) *Upper* – graphic comparing B6 vs. KI strains sharing the same genetic background and differing only in *Prdm9* allele. Paired horizontal bars indicate homologous chromosomes, height of peaks suggest level of open chromatin at hotspot. *Lower* – Normalized (cpm) profiles of histone modifications and chromatin accessibility at two hotspots activated by different *Prdm9* alleles. (B) MA plots comparing ATAC-seq (*upper*) and H3K9ac (*lower*) between B6 and KI. *Prdm9^Dom2^* (blue) and *Prdm9^Cst^* (red) hotspot annotations from (Baker et al. 2014). (C) *Upper* – B6 and D2 strains represent different genetic backgrounds sharing the same *Prdm9^Dom2^* allele. *Lower* – Heat maps showing H3K4me3, ATAC, SNV location, and PRDM9 motif scores at 1,331 local-QTL (Baker et al., 2018). (D) *Upper* – graphic outlining genetic makeup of heterozygous (B6xCAST)F1 hybrids with two *Prdm9* alleles. *Lower* – allele-specific profiles of H3K4me3, H3K9ac, ATAC, and PRDM9 (reads uniquely mapped to either the B6 or CAST haplotype). (E) Scatter plots of haplotype-specific PRDM9 binding (n = 3,786) versus H3K9ac (*left*) or ATAC (*right*). (F) *Upper* – open chromatin at hotspots occurs prior to programmed DSBs. *Lower* – H3K4me3, H3K9ac, and ATAC profiles in *Spo11^-/-^*mutant and H3K4me3 and ATAC profiles in heterozygous control at a representative hotspot (Pbx1). (G) Meta-profile of H3K4me3 and ATAC (*upper*) and H3K4me3 and H3K9ac (*lower*) at recombination hotspots from Fig. 1D from *Spo11^-/-^*. (H) MA-plot of ATAC-seq from *Spo11^-/-^*(mut, n = 2) and *Spo11^+/-^* (het; n = 1).

Reciprocally, we found 1,756 ATAC peaks and 7,049 H3K9ac peaks with increased levels in B6 compared to KI, with 94% of the ATAC peaks (n = 1,643) and 91% of the H3K9ac peaks (n = 6,431) overlapping known *Prdm9^Dom2^*-dependent H3K4me3 peaks. Similar to the case with B6, open chromatin at hotspots in the KI strain is highest at PRDM9 binding sites (**Fig. S1G** and **H**). These data show that hotspot chromatin that is accessible in B6 is not accessible in the KI. Instead, in the KI there are unique sites of accessible chromatin that overlap with known *Prdm9^Cst^*hotspots, sites that are not open in B6.

Residence time of DNA-binding proteins can be estimated *in vivo* using nuclease protection assays such as ATAC-seq and DNAse hypersensitivity (Sung et al., 2014). A large decrease in frequency of nuclease cutting at binding sites correlates with long residence times. *In vitro*, PRDM9 has a long residency time when bound to synthetic DNA oligonucleotides (Striedner et al., 2017). Footprint analysis of ATAC-seq data from germ cells found that PRDM9^Cst^ motif locations show little evidence for nuclease protection (**Fig. S1I**), relative to long-resident proteins like CTCF (**Fig. S1J** and **K**). The lack of a clear footprint is consistent with PRDM9 having a short residency time *in vivo*, similar to other transiently binding pioneer factors (Swinstead et al., 2016a).

Next, to determine if open chromatin at hotspots is PRDM9-dependent, we reasoned that genetic variants that disrupt PRDM9-binding should change accessibility. B6 and DBA/2J (D2) strains share the *Prdm9^Dom2^* allele, thus providing a natural experiment to determine the effect of genetic variation on chromatin accessibility. Recently we mapped quantitative trait loci (QTL) that regulate H3K4me3 levels in a genetic reference population whose genomes are homozygous mosaics of B6 and D2 (Baker et al., 2018). That study identified 1,331 hotspots where the H3K4me3 level is regulated by local-QTL. We performed ATAC-seq on D2 mice and compared H3K4me3 levels and open chromatin at local-QTL to that from B6 (**Fig. 2C**). Hotspots with biased H3K4me3 levels also had biased open chromatin. Local-QTL with the highest level of haplotype bias in ATAC signal showed an increased frequency of single nucleotide variants near PRDM9 binding sites. At each hotspot with a local-QTL we separately calculated scores for the PRDM9 motif (Baker et al., 2014) for both B6 and D2 sequences. Generally we found that the strain with the highest motif scores had the highest chromatin accessibility (**Fig. 2C**, right).

The programmed DSBs in meiosis are repaired primarily using the homologous chromatid as a template. This enabled us to determine whether the unique epigenomic landscape at hotspots occurs on both chromatids or only on the initiating chromatid. We performed ATAC-seq and H3K9ac ChIP-seq on spermatocytes from (B6xCAST)F1 hybrids and compared them to our published PRDM9 and H3K4me3 ChIP-seq data (Baker et al., 2015a). At asymmetric hotspots, i.e. those sites where PRDM9 preferentially binds one parental chromatid (Baker et al., 2015a; Davies et al., 2016; Smagulova et al., 2016), both acetylation and chromatin accessibility are largely restricted to the parental chromatid to which PRDM9 binds (Fig. 2D and E). These data demonstrate that open chromatin is concordant with PRDM9 binding. Interestingly however, they also suggest that hotspot chromatin is not open on the homologous chromosome, implying another mechanism is required to facilitate DNA accessibility for repair.

### Formation of open chromatin at hotspots precedes meiotic DSBs

In somatic cells, introduction of DSBs causes nucleosome remodeling and increased chromatin accessibility (Price and D’Andrea, 2013). To determine if this is also true of meiotic DSBs or if DSB formation follows chromatin remodeling, we tested if the epigenomic state at hotspots is dependent on DSB formation. We performed ATAC-seq and ChIP-seq for H3K9ac and H3K4me3 ChIP-seq on spermatocytes from mice lacking SPO11 (Baudat et al., 2000; Romanienko and Camerini-Otero, 2000). In the absence of SPO11, we found wild-type levels of open chromatin and H3K9ac that show no quantitative differences compared to heterozygous littermate controls (Fig. 2F-H). These data agree with previous observations showing that H3K4me3 and H3K36me3 modifications at hotspots are independent of and therefore precede SPO11 binding and subsequent DSB formation (Buard et al., 2009; Grey et al., 2017).

### The chromatin remodeling factor, HELLS, is required for proper synapsis

Our observations that hotspots are marked by histone modifications associated with active chromatin and show PRDM9-dependent formation of accessible chromatin all point to an unknown mechanism controlling chromatin remodeling in recombination. To identify candidate remodeling complexes, we reasoned that if chromatin remodeling is required for hotspot activation, loss of the remodeling enzyme would result in a phenotype similar to loss of *Prdm9*, including incomplete synapsis, persistent DSBs, and a block in early pachytene (Hayashi et al., 2005). Loss of either *Ino80* (Serber et al., 2016) or *Hells* (Zeng et al., 2011) fulfill these criteria. We focused our attention on *Hells* based on two observations. First, *Ino80* plays a major role in histone variant exchange, specifically H2A.Z and H2A (Papamichos-Chronakis et al., 2011), and recombination hotspots lack H2A.Z (**Fig. 1A**). Second, single-cell transcript analysis found that individual cells that co-express *Hells* and *Prdm9* are restricted to leptotene; in contrast, *Ino80* is expressed more broadly during meiosis (**Fig. S2**) (Jung et al., 2019).

To examine HELLS protein localization during spermatogenesis, we used immunohistochemistry to detect protein abundance in seminiferous tubules (**Fig. 3A**). HELLS shows high expression in both spermatogonia and meiotic cells, similar to a previous result (Zeng et al., 2011). In meiosis, HELLS expression overlaps with phosphorylated H2AX (γH2AX), a marker of DSBs, with broad temporal colocalization in leptotene and zygotene stages. During pachytene, both HELLS and γH2AX are sequestered to the sex body (**Fig. 3A**, arrow). HELLS is also co-expressed in leptotene with PRDM9 (**Fig. 3A**). Together, these data show that HELLS is present with PRDM9 at the stage when recombination hotspots are activated.

**Figure 3.**
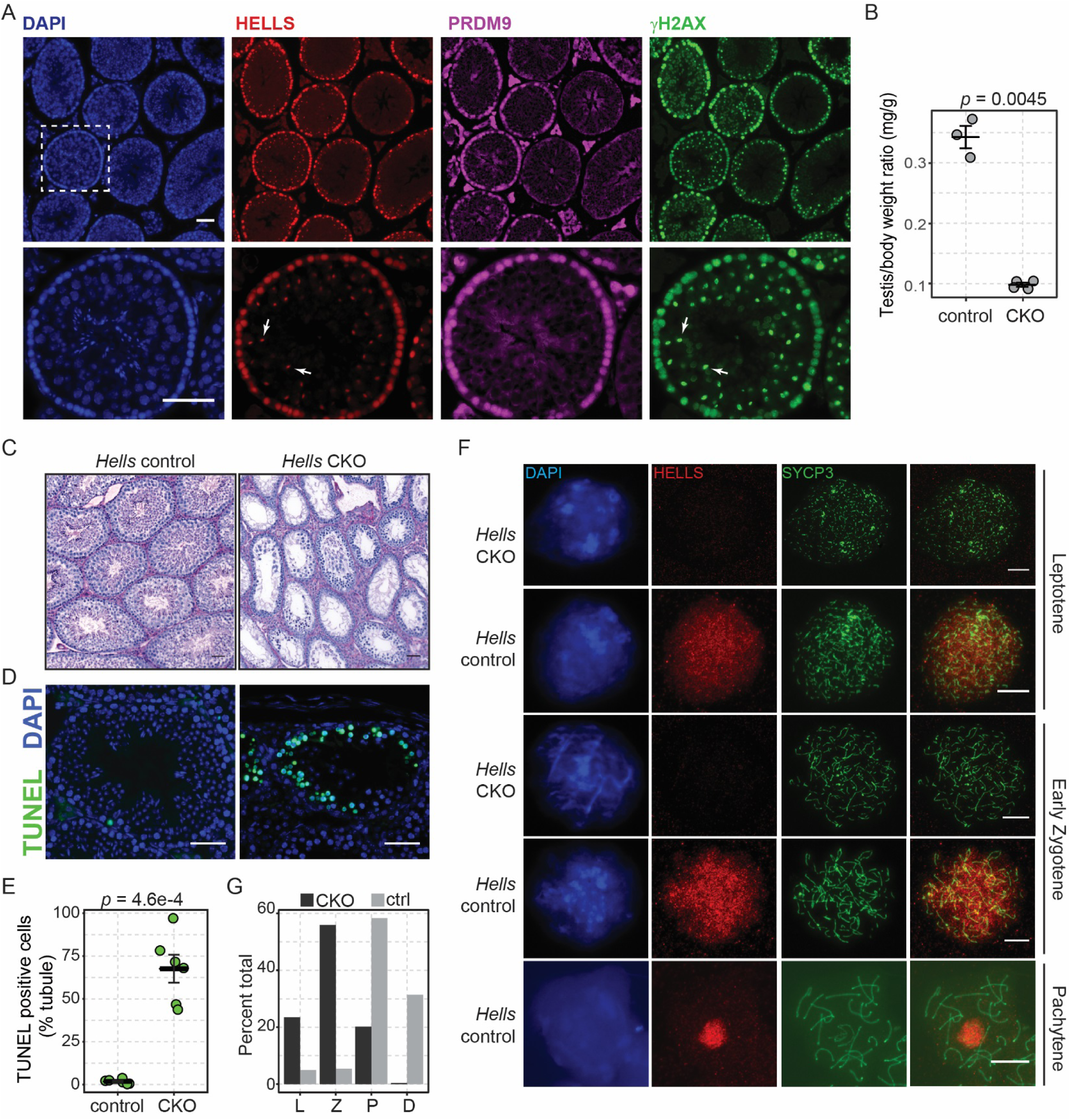
HELLS co-localizes with PRDM9 and is required for meiosis and spermatogenesis. (A) B6 adult testis cross sections immunolabeled with anti-HELLS (red), anti-PRDM9 (magenta), and anti-γH2AX (green). Nuclei were counterstained with DAPI in blue (arrow – sex body) (B) Testis weight compared to body weight in adult *Hells* control and CKO mice (mean ± SEM, *p* value – Welch’s two-sided test). (C) Adult testis cross sections stained with PAS. (D) TUNEL staining (green) of cross-sections of adult testis from *Hells* control and CKO. Nuclei were counterstained with DAPI in blue. *A*, *C*, and *D* scale bar = 50 µm. (E) Percent TUNEL positive cells per tubule (mean ± SEM, *p* value – Welch’s two-sided test). (F) Chromatin spreads of spermatocytes from *Hells* control and CKO animals stained for DAPI (blue) and immunolabeled with anti-HELLS (red) and anti-SYCP3 (green). Scale bar = 10 µm. (G) Quantification of meiotic prophase I stages, based on SYCP3 and γH2AX staining, in control and CKO mice (n = 200 cells from two mice per genotype).

To study the meiosis-specific function of HELLS we used a conditional strategy to generate mice that lack *Hells* specifically in male germ cells at the onset of meiosis. Adult male *Hells* homozygous conditional knock-out mice (*Hells* CKO) showed reduced testis weight (**Fig. 3B**) and extensive loss of germ cells beyond prophase I (**Fig. 3C**), while *Hells* heterozygous littermates (control) appear normal. This meiotic prophase I block recapitulates an earlier observation using allographs of testis tissue between *Hells^-/-^* mice and wild-type donors (Zeng et al., 2011). Cross sections of adult *Hells* CKO seminiferous tubules show loss of HELLS staining (**Fig. S3A**). The loss of later stage germ cells correlates with a large increase in germ cell apoptosis in HELLS CKO compared to heterozygous littermate controls (Fig. 3C and D). Finally, *Hells* CKO male mice did not sire any offspring when mated to normal B6 females. In summary, we confirm that HELLS is required for sperm production and fertility of male mice.

Using surface-spread meiotic chromatin in combination with immunohistochemistry, we found that HELLS is diffusely localized in leptotene and zygotene spermatocytes (**Fig. 3F**), similar to localization of PRDM9 in the same stages (Parvanov et al., 2017). Meiotic spreads showed that later in prophase I, after repair of the majority of DSBs during the pachytene stage, HELLS is restricted to the sex body. Staining for HELLS is absent from *Hells* CKO mice (**Fig. 3F**). To determine at which stage meiotic progression is blocked in *Hells* CKO mice, we staged spermatocytes using spreads stained for both γH2AX as a marker of DSBs and SYCP3 to characterize chromosome synapsis in adult male mice (**Fig. S3B**). We found that *Hells* CKO mice have an increased proportion of spermatocytes in leptotene and zygotene stages, and a nearly complete loss of normal pachytene and diplotene stages (**Fig. 3F**). *Hells* CKO cells scored as pachytene-like consistently show persistent γH2AX phosphorylation absent from littermate controls and increased incomplete synapsis (**Fig. S3B**). Overall, our data confirm that loss of *Hells* leads to meiotic arrest at the late-zygotene to early-pachytene stage (Zeng et al., 2011), with incomplete synapsis and persistent DSBs leading to spermatocyte apoptosis, a phenotype identical to that due to loss of *Prdm9* (Hayashi et al., 2005).

### In *Hells* null mice meiotic DSBs are targeted to other functional elements

Because loss of *Hells* leads to incomplete synapsis and persistent γH2AX, we next determined the levels and locations of meiotic DSB in the mutant spermatocytes. To characterize DSBs cytologically, we used spread meiotic chromatin labeled with SYCP3 and DMC1. DMC1 is a meiosis-specific recombinase that binds to single-stranded DNA at the sites of DSBs (Neale and Keeney, 2006). While zygotene spermatocytes from *Hells* CKO overall had similar levels of DMC1 foci as littermate controls (**Fig. 4A**), these foci persist in *Hells* CKO pachytene-like cells on synapsed chromosomes (**Fig. 4B**), indicating incomplete DSB repair.

**Figure 4.**
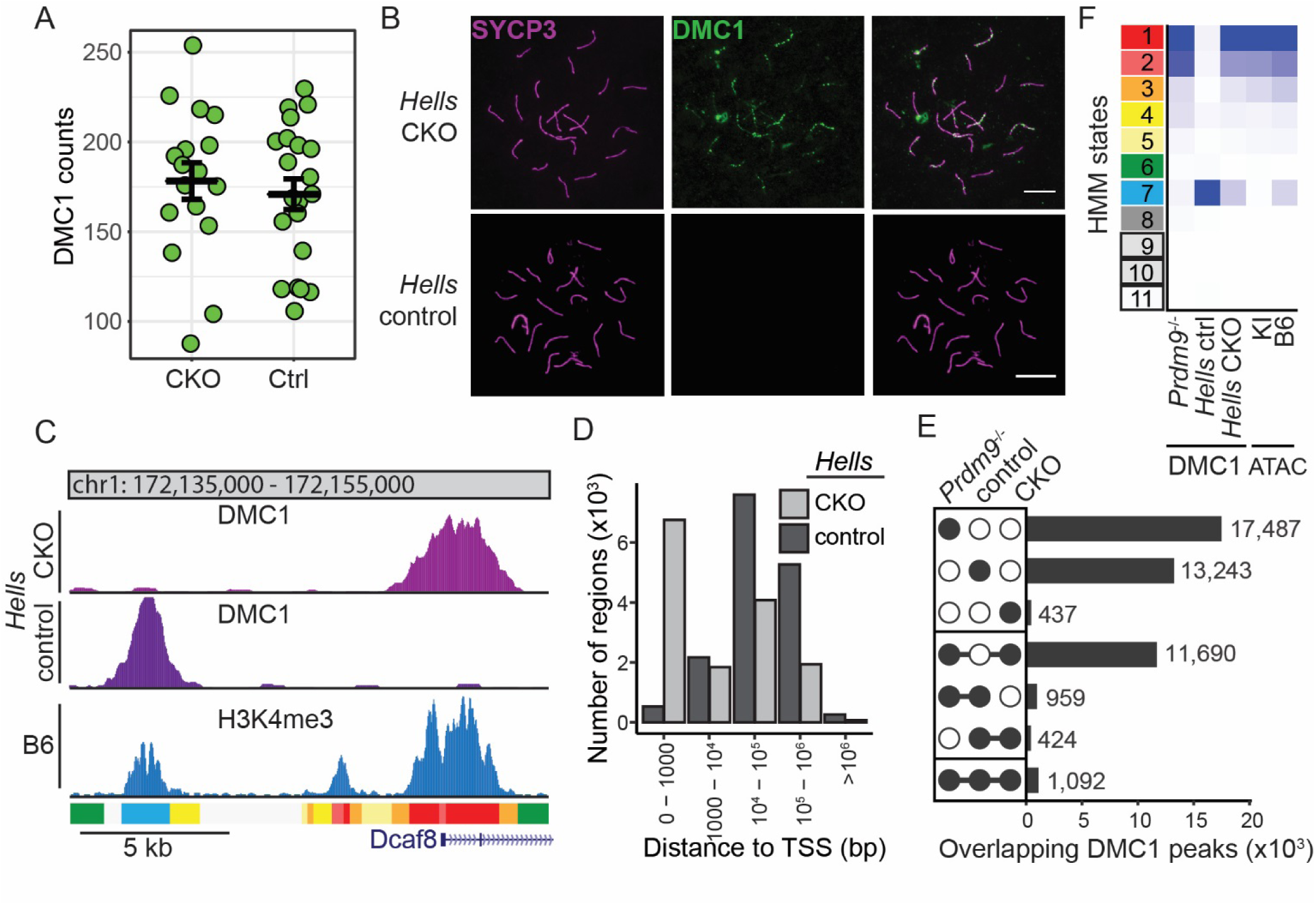
Loss of HELLS leads to redirection of programmed DSBs to other functional elements. (A) Quantification of DMC1 foci from chromatin spreads at zygotene/early pachytene stage spermatocytes in *Hells* CKO and control (Ctrl) animals (mean ± SEM). (B) Representative chromatin spreads of a pachytene-like spermatocyte from *Hells* CKO and pachytene spermatocyte from *Hells* control. Spreads were immunolabeled with anti-SYCP3 (magenta) and anti-DMC1 (green). Scale bar = 10 µm. (C) Profiles of DMC1 ssDNA signal and H3K4me3 level for representative locus in *Hells* CKO and control spermatocytes. ChromHMM state annotations are shown below profile tracks. (D) Distribution of DMC1 ssDNA regions relative to transcription start sites (TSS). (E) Intersection of DMC1 locations identified in spermatocytes from *Prdm9^-/-^* and *Hells* control and CKO mice. (F) Heat map showing fold enrichment of DMC1 locations and ATAC in ChromHMM states from Fig. 1A.

While the number of DSBs were similar between *Hells* CKO and control, their genomic locations were dramatically different. To determine locations of meiotic DSBs we used single-strand sequencing of DNA bound by DMC1 (Khil et al., 2012; Smagulova et al., 2011). In total, we identified 13,643 DMC1 peaks in *Hells* CKO and 15,718 in heterozygous littermates. Overall, *Hells* CKO germ cells lost DSBs at canonical hotspots and gained DSBs at other sites (**Fig. 4C**). For example, loss of *Hells* resulted in a 12-fold increase in DSBs at promoters, and a concomitant decrease in DSBs at distal sites (**Fig. 4D**). Nearly all DMC1 peaks identified in *Hells* CKO germ cells overlap with the location of DSBs in *Prdm9^-/-^* (93.7%, (13,206/13,643) (Smagulova et al., 2016); while only 15.7% (2,475/15,718) of DMC1 sites identified in control germ cells overlap *Prdm9^-/-^* DMC1 peaks (**Fig. 4E**).

To determine the epigenomic landscape targeted for DSBs in *Hells* CKO, we calculated enrichment of DMC1 sites within the ChromHMM states (**Fig. 4F**). While hotspots (state 7) were enriched for DSBs in the *Hells* control, in both *Hells* and *Prdm9* mutants DSBs were enriched in states 1 and 2, which are annotated as promoters, insulators, and enhancers. Interestingly, although H3K4me1, H3K4me3, and H3K9ac are present in states 1 and 2 (**Fig. 1A**), state 3 had a combination of histone modifications similar to classically defined enhancers that lack H3K4me3, and was less enriched for DSBs. States 1 and 2 were enriched for PRDM9-*independent* accessible chromatin; whereas state 7 only had PRDM9-*dependent* accessible chromatin in B6 compared to KI (**Fig. 4F**). Notably, unlike hotspots, states 1 and 2 were not enriched for H3K36me3. Furthermore, in *Hells* CKO, where DSBs largely occur at state 1 and 2 locations, spermatocytes undergo apoptosis similar to loss of *Prdm9* (Smagulova et al., 2011). Together, these data suggest that in the absence of *Hells*, DSBs are retargeted to regions of open chromatin which lack the proper epigenome to become competent hotspots.

### HELLS is required for epigenomic activation of recombination hotspots

Because DSBs were retargeted in *Hells* CKO germ cells, we next examined the epigenome of the mutant spermatocytes. After birth, the first wave of meiosis occurs semi-synchronously (Ball et al., 2016; Bellve et al., 1977), allowing enrichment of meiotic stages prior to *Hells* CKO meiotic arrest. We first confirmed loss of HELLS in 12 dpp whole testis (**Fig. 5A**). ChIP-seq for H3K4me3 and ATAC-seq on enriched germ cell populations from 12 dpp male mice identified a significant reduction of epigenomic modification in *Hells* CKO mice at recombination hotspots (Fig. 5B and C, Fig. S4). While 11,812 H3K4me3-modified sites are significantly reduced in *Hells* CKO compared to control (log_2_ fold change < −1, FDR < 0.01), only 73 sites showed higher modification in *Hells* CKO. Annotating peaks using state 7, we found that both H3K4me3 (n = 11,125, **Fig. 5D**) and chromatin accessibility (n = 1,851, **Fig. 5E**) sites at hotspots were reduced in *Hells* CKO. In contrast, among peaks overlapping HMM state 1 (n = 31,161 H3K4me3 peaks) there were virtually no sites that changed. Histological sections of 12 dpp seminiferous tubules from both *Hells* CKO and heterozygous control littermates showed similar stages in development (**Fig. S5A**). Although measuring PRDM9 in whole testis from CKO mice by western blot detected a modest reduction in protein abundance (**Fig. S5B** and **C**), PRDM9 was readily detected in individual leptotene and zygotene spermatocytes (**Fig. S5D**). These data show that HELLS is required for establishment of the epigenomic state and chromatin accessibility at hotspots.

**Figure 5.**
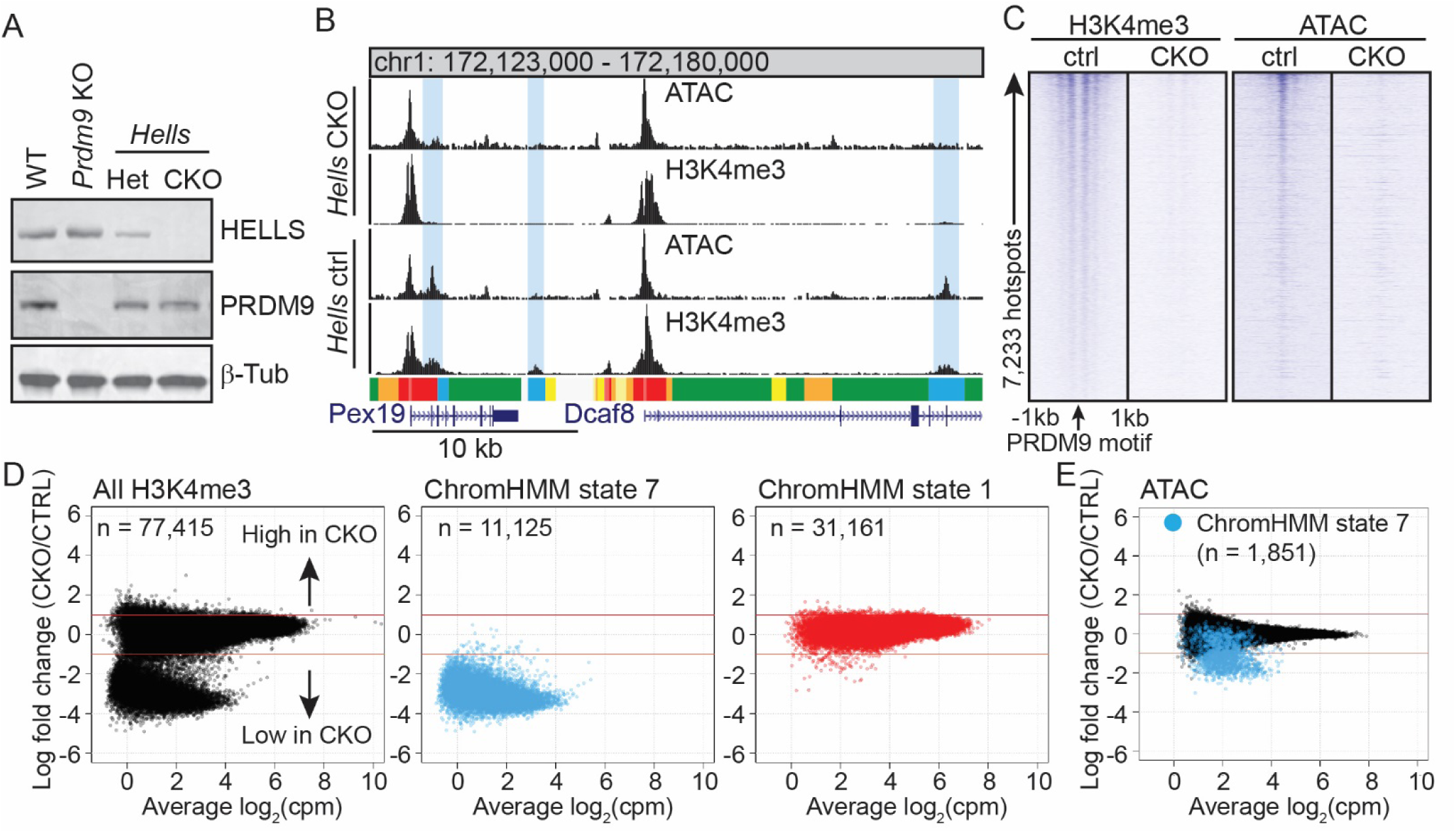
HELLS is required for histone modification and chromatin accessibility specifically at hotspots. (A) Western blot of whole testis protein extract from B6 (WT), *Prdm9^-/-^* (KO), and *Hells* heterozygous (Het) and CKO (β-Tubulin – loading control). (B) Profile of ATAC and H3K4me3 levels from 12 dpp *Hells* CKO and control spermatocytes. HMM annotations are below. (C) Heat map of H3K4me3 and ATAC signal at all hotspots from Fig 1D (cpm in 10 bp bins). (D) MA-plots of H3K4me3 levels comparing *Hells* CKO (n = 3) and homozygous controls (n = 3) for all H3K4me3 sites (*left*), HMM state 7 (*middle*, filtered for overlap with DSBs and PRDM9-dependent H3K4me3), and HMM state 1 (*right*). (E) MA-plot of ATAC signal (blue – HMM state 7, n = 2 control and n = 3 CKO).

### HELLS forms a complex with PRDM9

Expression of PRDM9 *ex vivo* recapitulates aspects of hotspot activation including allele-specific H3K4me3 deposition (Altemose et al., 2017; Baker et al., 2015b; Thibault-Sennett et al., 2018). To develop a model for temporal molecular characterization of hotspot activation, we choose a human cell line that expresses HELLS and created a stably-integrated FLAG-*PRDM9^C^* allele under inducible control of the tetracycline promoter (HEK293-P9^C^, **Fig. 6A**). Addition of doxycycline led to robust expression of FLAG-PRDM9^C^ and increased H3K4me3 at PRDM9^C^ hotspots (Fig. 6A and B). PRDM9^C^ expression also increased MNase sensitivity at PRDM9^C^ binding sites compared to adjacent regions (**Fig. 6C**), indicating nucleosome remodeling (Baker et al., 2014). Reciprocal immunoprecipitation using antibodies directed to either HELLS or PRDM9 (FLAG) identified a doxycycline-dependent interaction between the two proteins (**Fig. 6A** right). We confirmed PRDM9-HELLS interaction *in vivo* by performing co-immunoprecipitation (co-IP) on protein lysates prepared from 12 dpp mouse testis. We detected strong interaction between HELLS and PRDM9 in testis from B6 mice and littermate controls but not in *Hells* CKO (**Fig. 6D**) or PRDM9^Δ/Δ^ (**Fig. 6E**). To test if PRDM9-HELLS interaction requires assembly on chromatin, co-IP was repeated using a mouse strain expressing a PRDM9 variant that lacks the DNA-binding zinc-finger domains (PRDM9^ΔZF^) (Parvanov et al., 2017). We found that HELLS did interact with PRDM9^ΔZF^ (**Fig. 6E**), suggesting that this interaction is independent of PRDM9-directed DNA binding.

**Figure 6.**
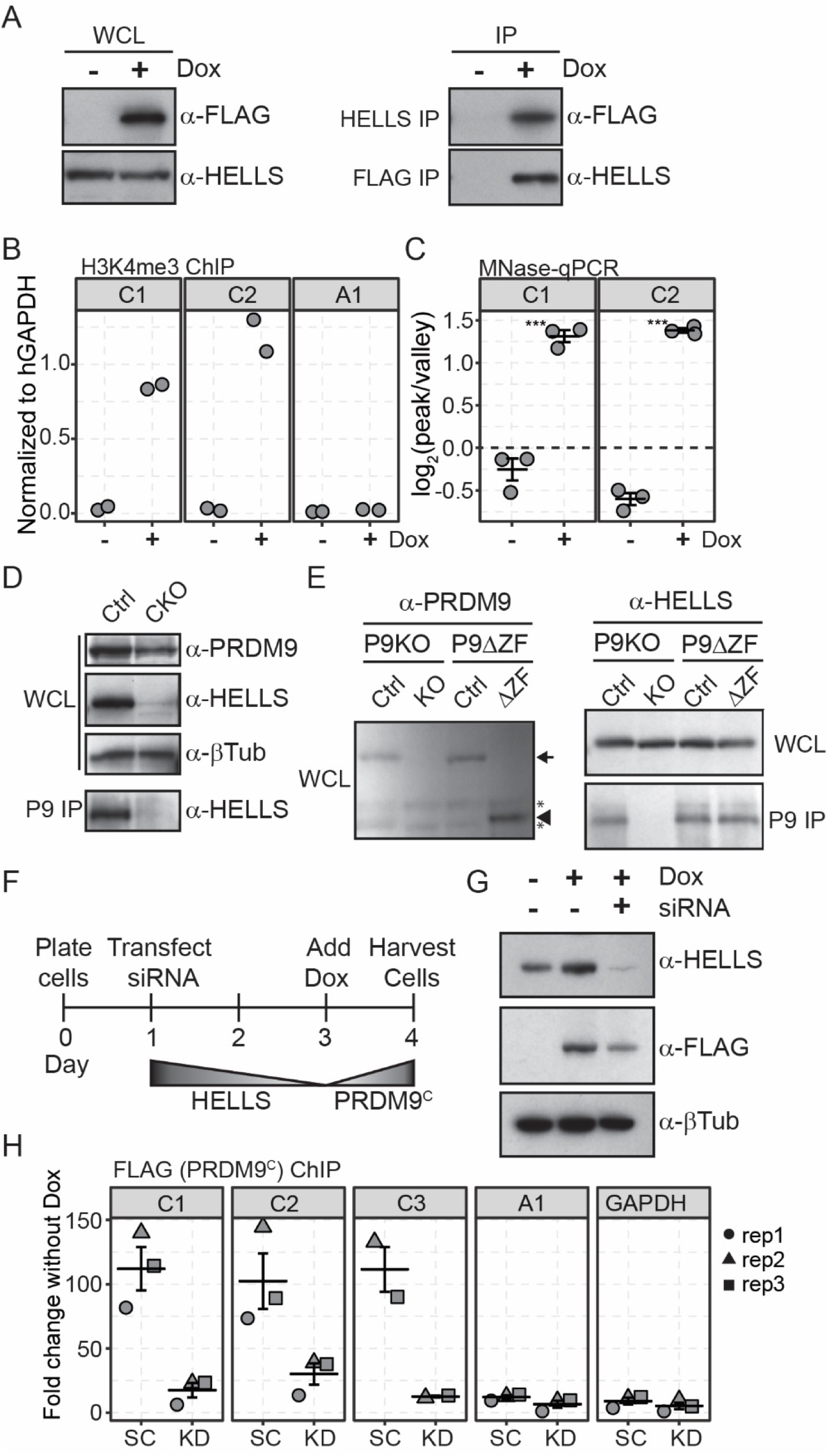
HELLS and PRDM9 form a complex *in vivo*. (A) Western blot of protein lysate from HEK293-P9^C^ cells without (-) or with (+) addition of doxycycline. (B) qPCR following H3K4me3 ChIP from HEK293-P9^C^ cells (C1 – Chr 3 144 Mb, C2 – Chr 3 4.4 Mb are PRDM9^C^ hotspots; A1 – Chr 3 55 Mb PRDM9^A^ hotspot). (C) qPCR of DNA following MNase-digestion of chromatin from HEK293-P9^C^ cells. Numbers represent ratio of CT values at the nucleosome peak vs. NDR (mean ± SEM, *** *p* values < 0.001; Welch’s two-sided test). (D) Western blots of either whole cell lysate (WCL) or PRDM9 IP (P9 IP) using protein lysate prepared from testis of either *Hells* control or CKO age-matched littermates. (E) Western blots of WCL and P9 IP from *Prdm9^-/-^* (P9KO) and *Prdm9^ΔZF^* (P9ΔZF) animals. Lysates were prepared from age-matched littermates (arrow – full-length P9, arrowhead – truncated P9, asterisk – non-specific). (F) Schematic of experimental design. (G) Western blots for WCL from HEK293 cells after addition of Dox, and siRNA for scrambled control (-) or *HELLS* (+). (H) qPCR of FLAG ChIP DNA following experimental design from *D*. Primers designed to PRDM9^C^ hotspots (C1-3), PRDM9^A^ hotspot (A1), or *GAPDH* promoter (mean ± SEM, SC – scrambled siRNA, KD – *HELLS* siRNA).

We next showed that HELLS is required for robust PRDM9 binding at hotspots (Fig. 6F-H). We first reduced HELLS expression in HEK293-P9^C^ cells using small interfering RNA (siRNA) and then induced expression of PRDM9^C^ (**Fig. 6G**). ChIP for FLAG-PRDM9^C^ in HEK293-P9^C^ cells, followed by qPCR, successfully detected robust binding of PRDM9^C^ at C-dependent hotspots and no binding at a PRDM9^A^ hotspot or at the GAPDH promoter (**Fig. 6H**). Critically, following siRNA knockdown of HELLS, PRDM9^C^ binding at C-hotspots was reduced to background levels (**Fig. 6H**). Together, these data show that PRDM9 and HELLS form a complex *in vivo* and, by extension, suggests that active chromatin remodeling is required for robust PRDM9 binding at hotspots.

### HELLS is recruited to recombination hotspots through PRDM9

Our experiments indicate that HELLS and PRDM9 interact and open chromatin at hotspots. This suggests a model in which PRDM9 recruits HELLS to hotspots to facilitate nucleosome remodeling for DSBs. To confirm this, we performed ChIP for HELLS.

We first tested whether HELLS binding at hotspots requires the presence of PRDM9 using our HEK293-P9^C^ cells (**Fig. 7A**). In the absence of doxycycline (no PRDM9^C^ expression), HELLS was not bound at any of the five genomic locations tested by qPCR. In contrast, upon expression of PRDM9^C^, we observed significantly increased binding of HELLS only at C-hotspots, and no change in binding at negative controls (**Fig. 7A**), suggesting HELLS binding at hotspots is PRDM9-dependent.

**Figure 7.**
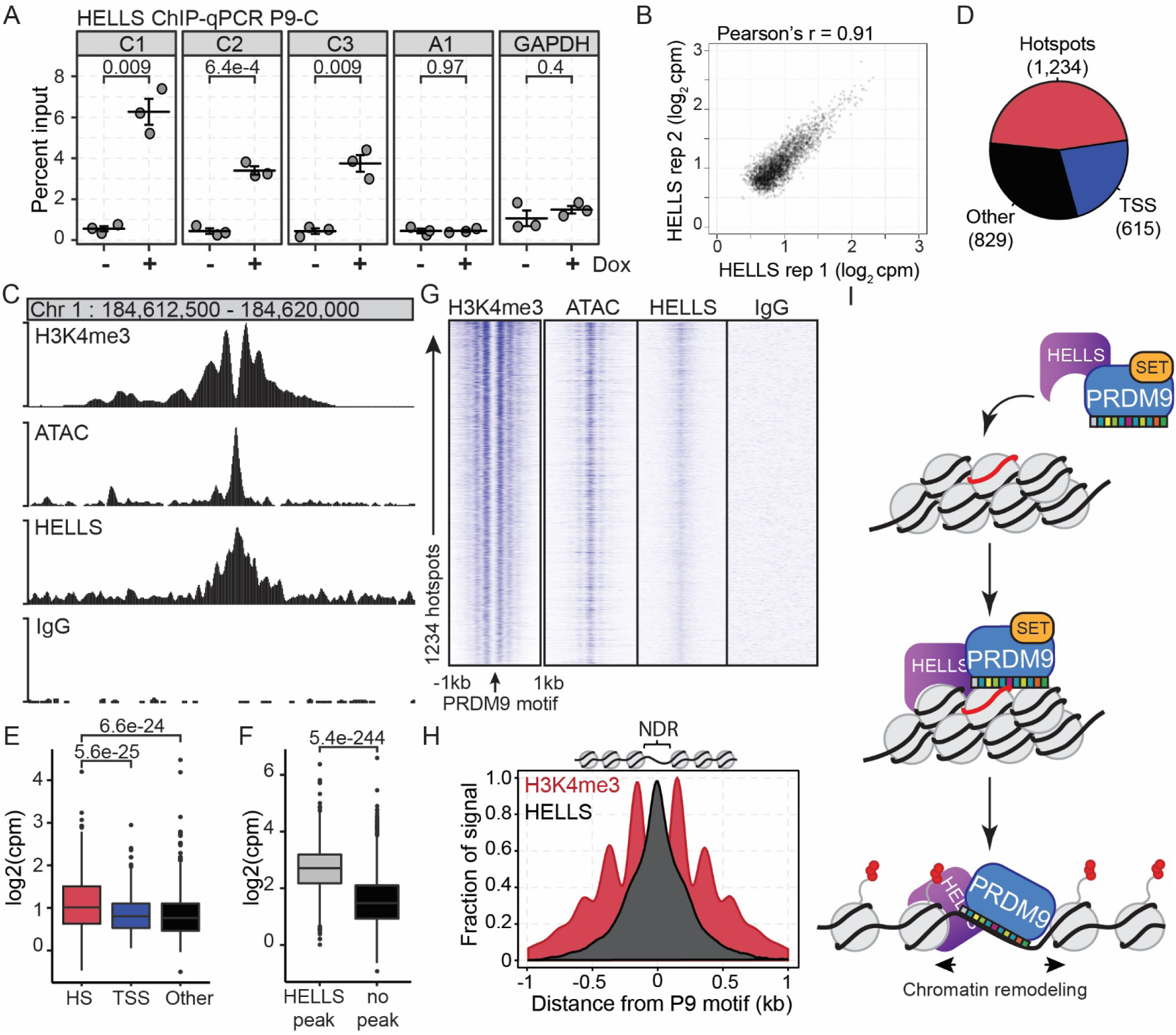
PRDM9 recruits HELLS to recombination hotspots. (A) qPCR of HELLS ChIP without (-) or with (+) addition of Dox in HEK293-P9^C^ cells (mean ± SEM; *p* values – Welch’s two-sided test). (B) Scatterplot of replicate HELLS ChIP-seq from mouse germ cell (n = 2,675 peaks) (C) Profile of H3K4me3, ATAC, HELLS ChIP-seq, and IgG control from germ cells of KI mice at the Hlx1 PRDM9^Cst^ hotspot. (D) Piechart annotating HELLS ChIP-seq peaks. (E) Boxplot comparing distribution of HELL binding (HS – hotspots/red; TSS – promoters/blue; Other – unknown elements/black; n = same as in *D*, *p* values – Welch’s two-sided test). (F) Boxplot comparing distribution of open chromatin at hotspots with (grey, n = 1,230) or without (black, n = 4,529) a HELLS ChIP-seq peak (*p* values – Welch’s two-sided test). (G) Heat maps of H3K4me3, ATAC, HELLS, and IgG signals +/- 1kb from PRDM9^Cst^ motif. (H) Meta-profile of H3K4me3 (red) and HELLS (grey) signals at hotspots from *G* (all HELLS signals are from merged replicates). (I) Model for PRDM9-dependent recruitment to recombination hotspots. PRDM9 and HELLS bind prior to interacting with chromatin. Nucleosome remodeling creates open chromatin surround PRDM9 binding site.

To test for HELLS binding of hotspots *in vivo*, we used enriched germ cells from 12 dpp KI mice, since the PRDM9^Cst^ allele reliably results in increased hotspot detection and higher signal-to-noise in functional genomic experiments (Baker et al., 2014; Grey et al., 2017) (**Fig. 2A** and **S2**). ChIP for HELLS in germ cells was highly correlated between biological replicates (Pearson’s r = 0.91, **Fig. 7B**) and identified 2,675 peaks. Visually, we detected clear HELLS ChIP signal at recombination hotspots (**Fig. 7C**) and promoters (**Fig. S6A)**. HELLS peaks were annotated based on overlap with PRDM9-dependent H3K4me3 peaks from KI mice (Baker et al., 2014) and TSS start sites representing promoters (**Fig. 7D**). The largest class of HELLS binding sites (n = 1,234, 46.1%) overlapped known PRDM9^Cst^ hotspots. As a class, HELLS binding was stronger at hotspots compared to promoters (p = 5.6e-25) or other unannotated regions (p = 6.6e-24, **Fig. 7E**). Of all HELLS binding regions, 97% (n = 2,591) fell within regions of open chromatin identified in KI germ cells. Moreover, hotspots with HELLS bound (n = 1,230) had significantly higher levels of open chromatin (p = 1.3e-244) compared to those without identified HELLS ChIP peaks (n = 4,529, **Fig. 7F**). In addition, genes with a HELLS peak at their promoters are generally expressed at significantly higher levels than those without (**Fig. S6B**). Given that HELLS is an ATP-dependent nucleosome remodeling factor, one model for HELLS interaction with chromatin would be recruited through histone modifications. However, the maximum average HELLS signal is found within the NDR at promoters (**Fig. S6C**) and at the PRDM9 binding site in hotspots (Fig. 7G-H). Together with the protein-protein interaction experiments, these results demonstrate that PRDM9 recruits HELLS to hotspots.

## DISCUSSION

Results here provide compelling evidence that HELLS and PRDM9 form a pioneer complex required for chromatin remodeling at recombination hotspots (**Fig. 7I**). Hotspots are marked by a unique epigenomic state not found at other functional elements, and are sites of accessible chromatin. This occurs exclusively on the chromosome PRDM9 binds and independently from DSBs. Furthermore, HELLS is required for proper targeting and repair of meiotic DSBs.

Pioneer function is defined by three features: 1) locational specificity; 2) active histone modifications; and 3) increased chromatin accessibility. Here we show that HELLS and PRDM9 fulfill these criteria at hotspots. PRDM9 brings two of the salient features of canonical pioneer factor-mediated chromatin reorganization into one molecule: 1) a DNA-targeting domain that provides locational specificity, and 2) the ability to catalyze epigenetic marks associated with active chromatin. Both of these features are required for meiotic recombination (Diagouraga et al., 2018; Parvanov et al., 2017). While the number of recombination hotspots are generally depleted from repressive regions of chromatin marked by H3K9me2/me3 (Patel et al., 2019; Walker et al., 2015), hotspots that are found in these domains show nearly similar level of PRDM9-dependent histone modification as those found outside of H3K9me2/3 domains, supporting the idea that even within heterochromatin, PRDM9 and HELLS have the capacity to create open chromatin. Further evidence for a pioneer-factor role for PRDM9 comes from our *ex vivo* analyses. Ectopic expression of PRDM9 in cells expressing HELLS recapitulates allele-specific histone modification and nucleosome remodeling. This suggests that the ability for PRDM9 to function as a pioneer factor is independent of chromatin structure that might be unique to meiosis.

Combined, our observations strongly support the hypothesis that the primary role of PRDM9 is to establish the proper open chromatin environment for both successful targeting of DSBs and DNA repair. The ability to target DSBs to open chromatin is critical for meiotic recombination. In mice, meiotic DSBs (Baker et al., 2014; Lange et al., 2016) and crossovers identified through single-sperm sequencing (Hinch et al., 2019) are concentrated within hotspot NDRs. In the absence of open chromatin at hotspots, by loss of either HELLS or PRDM9 (Brick et al., 2012), SPO11 preferentially targets other regions of accessible chromatin to create DSBs, even when the effect of these ectopic breaks is usually blocked meiotic progress and infertility. Notably, however, the severity of the infertility phenotype caused by loss of PRDM9 depends on the genetic background (Mihola et al., 2019), suggesting other components in action. Finally, through creation of aberrant sites of open chromatin, the pioneer function of the HELLS-PRDM9 complex could provide a molecular mechanism by which particular alleles are associated with genomic instability in certain cancers (Houle et al., 2018).

Here we have identified the chromatin remodeling enzyme HELLS is a second critical component of recombination initiation. Chromatin accessibility and histone modification at hotspots requires HELLS, and furthermore, HELLS localizes at hotspots through interaction with PRDM9. HELLS is a member of the SNF2 family of ATP-dependent chromatin remodeling enzymes that use the energy of ATP to reposition, remodel, and remove histones from DNA substrate (Clapier and Cairns, 2009). *HELLS* is misregulated in human tumors (von Eyss et al., 2012) and mutated in patients with immunodeficiency-centromeric instability-facial anomalies syndrome (Thijssen et al., 2015). Mice with systemic loss of HELLS exhibit prenatal lethality (Geiman et al., 2001); therefore, most functional studies of HELLS have required *ex vivo* cell culture. Notable exceptions are two studies showing that loss of HELLS in both male and female germ cells results in errors in synapsis and DSB repair during meiosis (De La Fuente et al., 2006; Zeng et al., 2011). Here, using an *in vivo* approach through conditional loss-of-function, we confirmed the requirement of HELLS in meiotic progression and defined the mechanistic role of HELLS in chromatin remodeling at hotspots.

HELLS has been implicated in the establishment of genome-wide methylation patterns and chromatin repression (Termanis et al., 2016; Yu et al., 2014). HELLS ATPase domain is required for these functions (Burrage et al., 2012; Ren et al., 2015; Termanis et al., 2016), and ATP is necessary for nucleosome remodeling *in vitro* (Jenness et al., 2018). If HELLS functioned to close chromatin in meiotic cells, we might have expected to see an increase in open chromatin or H3K4me3 level in CKO mice; instead we detected a near universal closing of chromatin at hotspots upon loss of HELLS. Given that our CKO strategy ablated HELLS primarily in meiotic cells, these epigenomic experiments are not optimal for determining the consequences of loss of HELLS in other cell types, but support the idea that the primary role of HELLS in meiosis is creating open chromatin specifically at hotspots. In contrast, ChIP for HELLS was performed on a population of male germ cells including spermatogonia. In addition to hotspots, we found that HELLS is enriched at promoters associated with higher gene expression, in agreement with previous results in fibroblasts (von Eyss et al., 2012). Together these data show that, in addition to its well-characterized role in generating repressive chromatin, HELLS is associated with increased chromatin accessibility. These disparate functions likely depend on protein binding partners.

Our evidence suggest that HELLS is recruited to hotspots through PRDM9, rather than being recruited as a consequence of PRDM9-dependent epigenomic modification. The HELLS-PRDM9 interaction persists in cells with a mutated PRDM9 lacking zinc fingers, supporting the idea that their interaction is independent of DNA binding by PRDM9. Second, the maximum HELLS ChIP enrichment is found in the center of NDR at the PRDM9 binding site, rather than at the flanking nucleosomes. These observations support a model where HELLS is recruited by direct interaction with PRDM9 and agree with the previous observation that HELLS-mediated chromatin remodeling *in vitro* requires interaction with the zinc-finger protein CDCA7 (Jenness et al., 2018).

Most pioneer factors contain a sequence-specific DNA binding domain that provides locational specificity, but the mechanistic details leading to the downstream opening of chromatin have remained elusive. The DNA-binding domains of canonical FoxA pioneer factors structurally resemble linker histones (Iwafuchi-Doi et al., 2016). This similarity first led to a physical model for chromatin opening whereby nucleosome packaging is disrupted through FOXA1 binding. However, most other pioneer factors, many of which are lineage-determining transcription factors and do not have DNA binding domains that resemble linker histones, are unlikely to open chromatin in this manner. Instead, it seems likely that sequence-specific DNA binding proteins need to recruit chromatin remodeling enzymes to open chromatin, such as the case with PRDM9 and HELLS. Supporting this hypothesis, another pioneer factor POU5F1 (OCT4) requires a chromatin remodeling enzyme, BRG1, in this case to create open chromatin in mouse embryonic stem cells (King and Klose, 2017).

Hotspots are marked by a unique combination of histone modifications not found at other regulatory elements, specifically H3K4me1, H3K4me3, H3K36me3, and H3K9ac. While PRDM9 is responsible for hotspot methylation, it is unknown which histone acetylase functions at hotspots. Candidates include proteins encoded by *Kat2a* (*Gcn5*) and *Kat2b* (*Pcaf*), known to catalyze acetylation at H3K9 (Jin et al., 2011). Specifically, KAT2A, which is part of the SAGA complex that is targeted to chromatin by prior H3K4me3 (Vermeulen et al., 2010). These observation support a precedent set by an early study that found enrichment of H3K9ac at an individual hotspot (Buard et al., 2009) and a recent survey of histone modifications in purified populations of male meiotic cells found that, out of 6 histone acetylation antibodies tested, only H3K9ac showed high enrichment at hotspots (Lam et al., 2019). It is possible that this unique combination of histone modifications provides an epigenomic addressing system for recruiting downstream DSBs and recombination factors to hotspots. One putative hotspot epigenomic reader could be ZCWPW1, which has domains predicted to bind H3K4me3 and H3K36me3 and is required for meiotic progression in males (Jung et al., 2019; Li et al., 2019).

Overall, data reported here give rise to a model (**Fig. 7I**) where HELLS and PRDM9 form a meiosis-specific pioneer complex to create open chromatin at recombination hotspots. This model not only expands our knowledge about diversity in function of pioneer complexes in general, but more specifically, also poses interesting challenges with respect to meiotic mechanisms. Clearly further work is required to determine how the HELLS-PRDM9 complex is temporally regulated, how the epigenomic environment and open chromatin at hotspots recruits programmed DSBs, and how chromatin barriers are overcome on the template chromatid used to repair these breaks.

## ACKNOWLEDGMENTS

We thank all members of the Baker laboratory for comments and discussion. We thank Anita Hawkins for assistance with obtaining mice with different *Prdm9* alleles, as well as Tanmoy Bhattacharyya and Travis Kent for technical help with PRDM9 histological staining and staging seminiferous tubules. We thank Mary Ann Handel and Taneli Helenius for critical reading of the manuscript and suggestions. This work was supported by P01 GM099640 to K.P and G.W.C., Jackson Laboratory start-up funds and R35 GM133724-01 to C.L.B., and was assisted by The Jackson Laboratory scientific services, which are supported through National Institutes of Health Cancer Core grant CA34196.

## AUTHOR CONTRIBUTIONS

Conceptualization, C.L.B.; Methodology, C.L.B. and G.W.C.; Formal Analysis, C.L.B. and G.A.; Investigation, C.S., S.D., N.B., H.T., and C.L.B.; Writing – Original Draft, C.S. and C.L.B.; Writing – Review & Editing, K.P. and C.L.B.; Funding Acquisition, K.P., G.W.C., and C.L.B., Supervision, C.L.B.

## DECLARATION OF INTERESTS

The authors declare no competing interests.

## METHODS

### Mice

C57BL/6J (B6, Stock No. 000664), DBA/2J (D2, 000671), CAST/EiJ (CAST, 000928), B6.129X1-*Spo11^tm1Mjn^*/J (*Spo11^-/-^*, 019117) (Baudat et al., 2000), and B6;129P2-*Prdm9^tm1Ymat^*/J (*Prdm9^KO^*, 010719) (Hayashi et al., 2005) are all available through The Jackson Laboratory catalog. Mice carrying various *Prdm9* alleles were previously described including B6.Cg-*Prdm9^tm1.1Kpgn^*/Kpgn (Baker et al., 2014) and B6.Cg-*Prdm9^tm3.1Kpgn^*/Kpgn (*Prdm9^ΔZnf^*, (Parvanov et al., 2017)) and are available upon request. For *Hells* loss-of-function, frozen embryos of the EUCOMM allele C57BL/6NTac-Hells^tm1a(EUCOMM)Wtsi^/Ieg were obtained through the European Mouse Mutant Archive (EMMA #04583) and implanted into pseudopregnant B6 females at The Jackson Laboratory. *Hells^tm1a(EUCOMM)Wtsi/+^* pups were crossed to B6.Cg-Tg(ACTFLPe)9205Dym/J (Stock No. 005703) to remove the *FRT* flanked neomycin selection cassette and make the conditional ready *Hells^tm1c(EUCOMM)Wtsi^*allele and resulting mice were subsequently inbred. Mice carrying *Hells^tm1c(EUCOMM)Wtsi^*conditionally ready allele are available through The Jackson Laboratory (B6.Cg-Hells^tm1c(EUCOMM)Wtsi^/Bakr, Stock No. 30483). Mice were made to carry a conditional homozygous germ cell-specific loss-of-function mice (CKO) in two steps by crossing *Hells^tm1c^* homozygotes to Tg(Stra8-iCre)1Reb/J (Stock No. 008208) (Sadate-Ngatchou et al., 2008) to create male mice with germ cells heterozygous for *Hells^+/tm1d^* and Stra8-iCre. These male mice were backcrossed to female *Hells^tm1c^* homozygotes to produce *Hells^tm1c/tm1d^*; Stra8-iCre offspring, of which testis from male mice would lack functional *Hells* (*Hells^tm1d^*homozygotes) and were used throughout this study as CKO. Fertility testing was performed on three Hells CKO male mice. Starting at 8 weeks of age, the male CKO mice were each mated to two or more confirmed breeder females for at least 4 months, and no pregnancy or litters were ever recorded. All animal experiments were approved by the Animal Care and Use Committee of The Jackson Laboratory (Animal Use Summary #16043).

### Cell culture and HEK293-P9^C^ cells

HEK293-P9^C^ cells were created using the T-REx^TM^ system (ThermoFisher #K102001) and the T-REx-293 (HEK293) cell line (ThermoFisher #R71007). Cloning of human FLAG-tagged PRDM9^C^ was previously described (Baker et al., 2015b), and subcloned into the doxycycline inducible expression system (pcDNA4/TO vector). T-REx-293 cells were transfected using lipofectamine 3000 (Invitrogen), allowed to recover for 48 hours, and subsequently split and plated on 100 cM plates to allow for clonal expansion and treated with 5 µg/mL blasticidin (ThermoFisher #R210-01) and 300 µg/µl Zeocin (ThermoFisher #R250-01) to select for integration of FLAG-PRDM9^C^ (HEK293-P9^C^). After 14 days individual colonies were picked, expanded, and tested by Western blot for expression of predicted full-length FLAG-PRDM9^C^. A single HEK293-P9^C^ cell line (#10) was expanded, tested for H3K4me3 deposition at hotspots, and subsequently used for all studies in this manuscript. All cell culture was performed using low glucose DMEM (Gibco 1885-084) and 10% FBS (Lonza 14-501F, lot#217266) at 37 °C at 5% CO_2_. FLAG-PRDM9^C^ expression was induced by replacing culture media with fresh media containing 1 µg/mL doxycycline (Sigma D9891) dissolved in DMSO (Sigma D2650-5×5ml) and grown for an additional 24 hours. Human *Hells* siRNA (Origene # SR302088) knock-down was performed in HEK293-P9^C^ cells with either 100 nM siRNA or universal scrambled control (Origene #SR30004) using Lipofectamine RNAiMAX following manufacture recommendations (ThermoFisher #13778030).

### Western blot, protein extraction, and immunoprecipitation

Protein extracts for Western blots from cultured HEK293-P9^C^ cells were prepared as previously described using RIPA buffer (Baker et al., 2015b). Protein extracts for co-immunoprecipitation from HEK293 cells was prepared using 200 µl HEPES-based buffer as previously described (Baker et al., 2015b). Whole testis were added to HEPES buffer (50 mM Hepes-KOH pH 7.4, 137 mM NaCl, 10% Glycerol, 0.4% NP-40) supplemented with 1 mM PMSF (Sigma 78830), 1X protease inhibitor cocktail (Sigma P8849), and 1 µl benzonuclease (Millipore/Sigma E1014-25KU) in 1.5 ml microtubes and homogenized using a micropestle on ice. Lysates were cleared by spinning at 10,000 x g for 10 minutes at 4 °C and supernatant transferred to a new tube. Cleared protein lysates were quantified using Bradford and compared against BSA standard. For IP, protein lysates were diluted to 500 µl using HEPES buffer and treated with 4 µl antibody and 20 µl Protein-A/G magnetic beads (Fisher). Immunocomplexes were incubated overnight at 4 °C with rotation, washed three times with HEPES wash buffer (50 mM Hepes-KOHpH 7.4, 150 mM NaCl, 0.4% NP-40), and eluted with 20 µl 2X sample loading buffer (50 mM Tris-HCL pH 6.8, 2% SDS, 10% Glycerol, 100 mM DTT, 0.05% Bromophenol Blue) heated at 95 °C for 10 minutes. Primary antibodies for western blots and immunoprecipitation include anti-FLAG M2 (Millipore/Sigma #F3165), anti-PRDM9 (custom (Parvanov et al., 2017)), anti-HELLS (Millipore/Sigma, ABD41 lot# 3069868), anti-β-Tubulin (Sigma, T4026 lot#125M4884V). Secondary antibodies for western blots include goat anti-mouse-HRP (BioRad 170-6516), goat anti-rabbit-HRP (BioRad172-1019), and donkey-anti-guinea pig (Millipore AP193P).

### Histological sections, chromatin spreads, and immunoflouresence

Testis from B6, *Hells* heterozygous control, or CKO mice were isolated from 6 week old and 12 dpp animals, fixed for 12-16 hours with Bouin’s solution, and embedded in paraffin wax. 5-µm sections were prepared and stained with Periodic acid-Schiff-diasase (PAS). Samples prepared for immunofluorescence were fixed in 4% PFA in PBS for 12-16 hours as previous described (Parvanov et al., 2017). Prior to immunofluorescent staining, sections were deparaffinized with xylene washes and rehydrated through a gradient of ethanol washes (95%, 70%, 50%, PBS) and then boiled in 10 mM sodium citrate buffer (pH 6.0) for 10 minutes. After being cooled and washed in PBS, the slides were blocked with 10% Normal donkey serum (017-000-121, Jackson ImmunoResearch Labs), 3% BSA, 0.2X protease inhibitor cocktail, and 0.05% Triton-X 100 for 1 hour at room temperature before adding primary antibodies and incubating overnight at 4°C. Slides were washed 3x 5 minutes in PBS and incubated in secondary antibody for up to 2 hours, then washed 3x 10 minutes in PBS. Slides were treated with ProLong Gold antifade reagent with DAPI (Invitrogen P36935).

Chromosome spreads were prepared using the drying-down technique (Peters et al., 1997). Testis tubules were lysed for 30 minutes in hypotonic extraction buffer (30mM Tris pH 8.2, 50mM sucrose, 17mM sodium citrate, 5mM EDTA, 2.5mM DTT, 1mM PMSF). After lysis, cells were liberated into 0.1M sucrose, then spread on a slide coated with 1% PFA. Slides were dried overnight in a humid chamber and rinsed in 0.04% Kodak Photo-Flo for 60 minutes before drying for immediate use or stored at −20°C. For immunofluorescence, slides were blocked with 3% BSA, 1% Normal donkey serum, and .005% Triton-X 100 between 1-6 hours before adding primary antibodies and incubating overnight. Slides were washed and treated with secondary antibodies as described above and treated with ProLong Gold antifade reagent with DAPI (Invitrogen P36935)

Primary antibodies used for Immunofluorescence on cross-sections and chromosome spreads include anti-PRDM9 (1:100, (Parvanov et al., 2017), anti-HELLS (1:250 Millipore/Sigma, ABD41), anti-γH2AX (1:2,000 Millipore 05636I, lot#2888552), anti-SYCP3 (1:500 Santa Cruz SC74569, lot#J1314), and anti-DMC1 (1:200 Santa Cruz, sc-8973). Secondary antibodies include: goat anti-rabbit immunoglobulin G (IgG; H+L), Alexa Fluor 594 conjugate (A-11037), 1:300; goat anti-rabbit IgG (H+L), Alexa Fluor 488 conjugate (ab150077), 1:500; goat anti-mouse IgG (H+L), Alexa Fluor 488 conjugate (ab150113); goat anti-mouse IgG (H+L), Alexa Fluor 647 conjugate (A-21236), 1:300; goat anti-guinea pig IgG (H+L), Alexa Fluor 488 conjugate (A-11073), 1:300. DMC1 spreads: donkey anti-goat IgG (H+L), Alexa Fluor 594 conjugate, 1:300, donkey anti-rabbit IgG (H+L), Alexa Fluor 488 conjugate, 1:500. Prdm9 sections: goat anti-guinea pig IgG (H+L), Alexa Fluor 488 conjugate (A-11073), 1:300, goat anti-rabbit immunoglobulin G (IgG; H+L), Alexa Fluor 594 conjugate (A-11037), 1:300, goat anti-mouse IgG (H+L), Alexa Fluor 647 conjugate (A-21236), 1:300. Sycp3 and γH2AX spreads: goat anti-rabbit immunoglobulin G (IgG; H+L), Alexa Fluor 594 conjugate (A-11037), 1:500, goat anti-mouse IgG (H+L), Alexa Fluor 488 conjugate (ab150113), 1:500.

TUNEL assay was performed on testis sections from 6 week old *Hells* heterozygous control or CKO mice using the In Situ Cell Death Detection Kit (Roche, #11684795910) according to manufacturer’s protocol.

### Chromatin immunoprecipitation and ChIP-seq library preparation

Isolation of enriched population of germ cells for ChIP for histone modifications and ATAC-seq was perform as previously described (Baker et al., 2014) on 12-14 dpp mice. Briefly, a single cell suspension was created using a combination of enzymatic and mechanical disaggregation. Seminiferous tubules were treated with 0.05 mg/ml Liberase TM (Roche) in 10 mM HEPES (pH 7.4) buffered DMEM with incubation at 37 °C for 15 minutes. Fragmented tubules were allowed to settle to separate interstitial cells from germ cells, and washed once with DMEM. Tubules were then resuspended in DMEM with Liberase and 500 µg/ml DNAse and incubated again at 37 °C for 15 minutes, disrupted with repeated pipetting, and passed through a 40 µM filter to remove cell clumps. This process removes the majority of somatic Sertoli and Leydig cells and results in over 90% enrichment of germ cells at different meiotic stages (Ball et al., 2016). Cells were subsequently crosslinked using 1% formaldehyde for 10 minutes and quenched using 125 µM glycine. Cells were hypotonically lysed, and nucleosome fragments solubilized by treatment with micrococcal nuclease. Chromatin was incubated for 2 hours at 4 °C with rotation with protein G magnetic Dynabeads preloaded with the indicated antibodies. Antibodies used for histone ChIP experiments were anti-H3K4me1 (Millipore/Sigma #07-436; lot: 2289129), anti-H3K4me3 (Millipore/Sigma #07-473; lot: 3018770), anti-H3K9ac (Active Motif #39137; lot:09811002), anti-H3K27ac (abcam #ab4729; lot:GR211893-1), anti-H3K27me3 (Millipore/Sigma #07-449; lot:2475696), anti-H2BK120ac (Active Motif #39120 lot:01008001), and anti-H2A.Z (Millipore/Sigma #07-594, lot: 2455725). For ChIP of H3K9ac, H3K27ac, and H2BK120ac used in the ChromHMM analysis, germ cell isolation and ChIP buffers were supplemented with 20 mM sodium buyterate to inhibit histone deacetylase activity. For H3K4me3 ChIP from *Hells* mice, one library was prepared using spermatocytes collected from 12-dpp *Hells^tm1c/+^* mice as control and three independently collected *Hells* CKO samples (**Table S1**, **Fig. S4**).

ChIP for FLAG-PRDM9 was previously described (Baker et al., 2015a) and modified for HELLS to include a dual crosslinking step to first capture protein-protein interactions. For HELLS ChIP from mice, tunica albuginea removed from testis and tubules were transferred to 5 ml of PBS pH 7.4 supplemented with 1 mM MgCl_2_. A fresh 0.25 M stock of disuccinimidyl glutarate (DSG) was prepared by dissolving in DMSO. DSG was added to tubules at a final concentration of 2 mM and incubated at room temperature for 30 minutes with rotation. After incubation, fresh paraformaldehyde was added directly to a final concentration of 1% and tubules were incubated for an additional 5 minutes with rotation, and quenched with 125 mM glycine. The remaining procedure for ChIP was followed as described (Baker et al., 2015a) using 4 µL anti-HELLS (Millipore/Sigma ABD41. lot# 3069868) or normal mouse IgG (Millipore/Sigma #12-371, lot: 2880788) per 3 million germ cells.

ChIP libraries were prepared for high-throughput sequencing either using Bioo Scientific’s NEXTflex ChIP-seq Kit without size selection (for ChromHMM data) or the KAPA hyper kit (Kapa Biosystems) for all other libraries. Library quality and size distribution were visualized using a Bioanalyzer (Agilent). All samples were sequenced using either the Illumina HiSeq2500 or NextSeq platforms.

Quantitative PCR on purified ChIP DNA was performed using PowerUp SYBR Green 2x Master Mix (ThermoFisher) on a ViiA 7 Real-Time PCR System (ThermoFisher). PCR was carried out for 40 cycles followed by melting curve analysis. All samples were run with triplicate technical reactions and cycle number was determined using an automated threshold analysis. MNase sensitivity assay in HEK293-P9^C^ cells before and after doxycycline addition was performed as described for spermatocytes (Baker et al., 2014). Oligonucleotides used for qPCR in this study are listed in **Table S2**.

### DMC1 ssDNA sample preparation and sequencing

Single-strand DMC1 ChIP-seq was performed as described (Khil et al., 2012) using DMC1 antibody (Santa Cruz sc-8973, lot: K2614) on testis from 6 week old male mice collected from two independent biological replicates of *Hells* CKO and heterozygous littermate controls (*Hells^tm1c/+^*; Stra8-iCre). Because of the reduced size of testis, three *Hells* CKO testes were used for each replicate, and only a single testis for each *Hells* control. Libraries were sequenced on Illumina HiSeq2500 platform with paired-end 75 bp reads.

### ATAC-seq sample preparation and sequencing

For B6, KI, D2, and (BxC)F1 hybrids genotypes ATAC-seq was performed on male germ cells using the original protocol (Buenrostro et al., 2013). ATAC-seq for *Spo11* and *Hells* genotypes was performed using the modified FAST-ATAC protocol (Corces et al., 2016). For *Hells* genotypes ATAC-seq libraries were prepared from two independent homozygous *Hells* wild-type (*Hells^tm1c/+^*) and three independent *Hells* CKO (*Hells^tm1c/Δ^*; Stra8-iCre). Tn5 transposition was performed using 1 x 10^5^ enriched male germ cell populations prepared from pooling testis from 3-4 individual mice similar to ChIP. ATAC-seq libraries were amplified by PCR using barcodes compatible with Illumina sequencing platform as described (Buenrostro et al., 2013) using a total of 8-10 amplification cycles and DNA purified using Agencourt AMPure XP beads (Beckman Coulter). Libraries were sequenced on either HiSeq2500 or NextSeq platforms either as paired-end or single-end libraries (**Table S1**).

### Data analysis

All reference sequence coordinates in the manuscript correspond to UCSC mm10 (release 3/18/2013).

Chromatin segmentation was performed using ChromHMM as described (Ernst and Kellis, 2012) solving for an 11-state solution. These data cover histone modifications commonly used to generate chromatin annotations in other cell types, including histone methylation associated with active chromatin (H3K4me3, H3K4me1), transcriptional elongation (H3K36me3), and repressed/inactive chromatin (H3K27me2, H3K9me2/3), as well as histone acetylation (H3K9ac, H3K27ac, H2BK120ac). We also performed ChIP-seq for the histone variant H2A.Z based on previous studies identifying an enrichment for this histone variant at recombination hotspots in plants (Choi et al., 2013) and fungi (Yamada et al., 2018). All datasets used for segmentation are outlined in **Table S1** with corresponding GEO accession numbers. Fold enrichment for chromatin states were calculated using transcription start sites (TSS), CpG islands, gene annotations, as well as previous published results for whole testis CTCF occupancy and predicted *cis*-regulatory elements (Shen et al., 2012), lamina-associated domains from mouse embryonic stem cells (Meuleman et al., 2013), *in vitro* PRDM9 binding sites (Walker et al., 2015), and gene expression levels from germ cells of age-matched mice (Ball et al., 2016). Duplicate ChIP libraries were merged prior to ChromHMM processing.

ATAC-seq fastq files were trimmed using trimmomatic (Bolger et al., 2014) using TRAILING:3 SLIDINGWINDOW:4:15 MINLEN:36 and removing Nextera adapters. For paired-end ATAC-seq samples, trimmed sequences were aligned to the mouse reference genome (mm10) using bowtie (version 0.12.9) (Langmead et al., 2009) with the following settings: -- chunkmbs 2000 -S -X2000 -m1. Mapped reads were adjusted for Tn5 insertion site as described (Buenrostro et al., 2013). For single-end ATAC-seq, trimmed sequences were aligned using bwa mem (version 0.7.15) (Li and Durbin, 2009) with default settings. Picard Tools was used to remove duplicates and calculate insert size metrics. ATAC-seq alignments were filtered to remove reads with MAPQ < 10 and mitochondrial reads.

ChIP libraries were aligned using bwa mem with default settings (version 0.7.15, (Li and Durbin, 2009)). In order to account for genetic variation, D2 ChIP-seq and ATAC-seq data were aligned to an *in silico* pseudogenome incorporating known variants (R78-REL1505) (Wu et al., 2010), and converted to mm10 reference coordinates using G2Gtools (accessible online at: https://github.com/churchill-lab/g2gtools). Allele-specific ChIP analysis was performed as previously described (Baker et al., 2015a) using variant-aware alignment strategy EMASE (Raghupathy et al., 2018). To identify hotspots, peaks were selected for those loci that overlap (BxC)F1 PRDM9 ChIP-seq summits. These were further filtered for peaks with greater than 30 reads that map uniquely to either B6 or CAST haplotype after removing duplicates to ensure robust haplotype calls.

Both ChIP-seq and ATAC-seq peaks were called using MACS version 1.4.2 (Zhang et al., 2008) using a cutoff of e-5. Duplicate reads were retained for ChIP against histone modifications and removing for HELLS ChIP. To build reference peakomes for each experiment, peaks were called on merged bam files from replicate experiments. Peak files across different strains or *Prdm9* alleles were concatenated, sorted, and merged using bedtools (Quinlan and Hall, 2010). Peaks overlapping ENCODE blacklist regions (Consortium, 2012) were removed for subsequent analysis. Read counts for each reference peakome were extracted from sample bam files using bedtools multicoverage and are available as processed data at NCBI GEO (GSE135896). Genomic profiles were visualized using the UCSC browser (Kent et al., 2002).

RNA expression data sets were from 14 dpp male germ cells downloaded from GEO GSE72833 (Ball et al., 2016). For ChromHMM annotation, expressed genes were filtered for transcripts per million (TPM) greater than 3 (n = 13,739). All remaining genes with Ensemble ids were annotated as ‘silent’. To visualize expression, two replicate 14-dpp RNA datasets (GSM1873271 and GSM1873272) were aligned to the reference genome (mm10) using bwa mem with default settings, merged, converted to bedgraphs using bedtools (Quinlan and Hall, 2010), and visualized using the UCSC browser.

All statistical analysis was performed using R (release 3.4.1). For differential analysis read counts were normalized using the trimmed mean of M-values (TMM) method, which accounts for sequencing depth and composition. Differential ATAC-seq peaks were identified using a generalized linear model which included batch in the model design comparing *Hells* CKO (n = 3) and homozygous controls (n = 2). Differential ChIP peaks were identified using the exact test function as implemented in edgeR (Robinson et al., 2010) comparing *Hells* CKO to *Hells* homozygous and B6 controls (**Fig. S4**). Genome-wide false discovery rates (FDR) were calculated by adjusting p-values following the Benjamini-Hochberg method. Annotation of peaks using genomic locus overlap was performed with bedtools intersect or HOMER mergePeaks functions (Heinz et al., 2013) and visualized using the UpSet package (Lex et al., 2014). Read matrices for heat maps were generated using the CoverageView R package after RPM normalization with 10 bp bins and visualized using Java TreeView (Saldanha, 2004). ATAC-seq footprints were aggregated and normalized using ATACseqQC (Ou et al., 2018). Calculation of motif scores for predicted PRDM9 binding sites was performed by generating a position weighted matrix for *Prdm9^Dom2^* motif (Baker et al., 2014) as outlined (Wasserman and Sandelin, 2004):

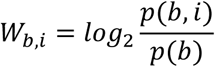

Where *Wb,i* = position weighted matrix value of base *b* in position *i*, where *p(b)* = the background probability of base *b* is set to 0.25. For each individual putative B6 and D2 binding sequence the *log2* values for each nucleotide in the PWM was summed to create an individual score.

