## Supplemental Material for "HELLS and PRDM9 form a Pioneer Complex to Open Chromatin at Meiotic Recombination Hotspots"

**Figure S1. Reproducibility of ATAC in germ cells.** (A) Scatterplot of ATAC-seq read counts from replicate B6 spermatocytes using 100,000 cells. (B) Distribution of insert size metrics for B6 ATAC-seq replicates from *A* showing clear nucleosome profile. (C) Enrichment of open chromatin around TSS sites from ATAC-seq replicates from *A*. (D) Scatterplot of ATAC-seq read counts from replicate KI spermatocytes using 100,000 cells. (E) Distribution of insert size metrics for KI ATAC-seq replicates from *D* and (B6xCAST)F1 showing clear nucleosome profile. (F) Similar to *C* showing TSS enrichment for KI and (B6xCAST)F1 samples. (G) Heat maps of H3K4me3, H3K9ac, and ATAC signals spanning ±1kb of previously identified PRDM9 motif locations. Heat maps are normalized reads per million (rpm) in 10 bp bins ordered by descending H3K4me3 level. Color scale is maintained across all panels. (H) Meta-profile of H3K4me3 level and open chromatin at recombination hotspots from KI spermatocytes anchored by PRDM9$^{Cst}$ motif (n = 9,305) scaled to maximum value to highlight positional relationship between nucleosome occupancy and open chromatin. (I) ATAC-seq cleavage footprint for PRDM9$^{Cst}$ show poor protection from Tn5 cutting across PRDM9$^{Cst}$ motif (n = 4,014 ATAC peaks that overlap PRDM9$^{Cst}$ motif) (J) Meta-profile of H3K4me3 (red) and ATAC (grey) signals at CTCF sites. CTCF ChIP-seq data is from modENCODE project (Shen et al., 2012). Only CTCF sites that did not overlap TSS were used (n = 2,926 ATAC peaks that overlap CTCF motif). Signals are background subtracted and scaled to maximum value. (K) Similar to *I* except for CTCF locations from *J* showing protection from cutting across CTCF motif locations (motif used in analysis shown above).

**Figure S2. *Hells* is expressed in spermatagonia and leptotene stages.** (A-F) t-SNE plots of single-cell RNA-seq from adult B6 testis (Jung et al., 2019) arranged by pseudo-time. All t-SNE plots were created using the testis Atlas shiny app (http://www.stats.ox.ac.uk/~wells/testisAtlas.html). (A) Highlighted cells are from cluster 5 annotated as leptotene stage. Arrow indicates pseudotime of spermatogenesis from spermatogonia to sperm. Individual meiotic stages are indicated by different colors. (B) t-SNE plot showing cells expressing *Prdm9* found largely in cluster 5 indicating leptotene stage. (C) t-SNE plot showing expression of *Hells* in spermatogonia and leptotene stages. (D) t-SNE plot showing *Ino80* expression across multiple stages of spermatogenesis.

**Figure S3. Meiotic-specific loss of *Hells* leads to zygotene/early-pachytene block.** (A) Immunolabeling of cross-sections from *Hells* CKO adult testis show loss of HELLS expression, persistence of DSBs in the most advanced meiotic cells, and lack of post-meiotic cells. Sections were immunolabeled with anti-HELLS (red) and anti-yH2AFX (green). Nuclei were counterstained with DAPI in blue. Scale bar = 100 µm. (B) Chromatin spreads of spermatocytes from *Hells* control and CKO animals displaying representative images of meiotic substages scored to determine meiotic block in **Fig. 3F**. Spreads were immunolabeled with anti-yH2AX (red) as an indicator of double-strand breaks and anti-SYCP3 (red) to allow visualization chromosome condensation. Scale bar = 10 µm.

**Figure S4. Loss of *Hells* results in decreased H3K4me3 at hotspots.** (A) Upper – dendrogram of hierarchical clustering (Euclidean distance) showing relationship among H3K4me3 ChIP-seq libraries for different mouse strains. Lower – boxplot comparing distribution of H3K4me3 level at hotspots (blue) compared to all other loci (grey). B6 12 dpp H3K4me3 ChIP-seq samples are from (Baker et al., 2014). (B) Scatterplot comparing single *Hells* CKO sample to age-matched B6 sample. Hotspots (blue) show higher H3K4me3 level in B6 compared to *Hells* CKO, while all other H3K4me3 sites (black) are more similar. Linear regression lines are plotted separately for both classes of H3K4me3 sites. (C) Scatter plot similar to *B* comparing two *Hells* CKO replicates. (D) Linear regression of pair-wise comparison between B6, *Hells* wild-type, and *Hells* CKO H3K4me3 ChIP samples. Linear regression lines are plotted separately for hotspots (blue) and all other H3K4me3 sites (black). Plot from panel *B* is highlighted in yellow and plot from panel *C* is highlighted in red.

**Figure S5. Testis from juvenile mice display similar stages of early meiotic progression in the absence of HELLS** (A) 12 dpp testis cross-sections, stained with PAS, of *Hells* heterozygous control and CKO testis. (B) Western blot of whole cell protein extract from whole testis (2 testis per genotype) collected from *Hells* homozygous controls, *Hells* CKO, and *Prdm9*$^{-/-}$ (P9KO). Western blots for *Hells* genotypes were collected from age-matched littermates and show two additional replicate Western blots in addition to **Figs. 5A and 6G**. Blots were decorated with anti-HELLS and anti-PRDM9, anti-β-Tubulin serves as a loading control. (C) Quantification of Western blots comparing PRDM9 expression levels between *Hells* CKO and age-matched littermate controls. Within each replicate the levels of PRDM9 in heterozygous control animals is set to one (replicate 1 is from **Fig. 5A**, replicate 4 is from **Fig. 6G**). β-tubulin was used as a loading control. (D) Immunolabeling of 12 dpp testis cross-sections of *Hells* heterozygous control and CKO testis show loss of HELLS expression in meiotic cells from CKO mice (arrow head – HELLS expression in pre-meiotic cells prior to Stra8-iCre expression, asterisk – incomplete loss of HELLS likely due to leaky *Cre* expression). In the absence of HELLS, PRDM9 expression is still detected along with DSBs. Sections were immunolabeled with anti-HELLS (red), anti-PRDM9 (magenta), and anti-P-H2AFX (green). Nuclei were counterstained with DAPI in blue. Scale bars = 50 µm

**Figure S6. HELLS binds at promoters.** (A) Profile of ATAC-seq, H3K4me3 ChIP-seq, IgG control, and HELLS ChIP-seq, at the *Sycp1* promoter in germ cells collected from PRDM9$^{Cst}$ hotspot in KI mice. RNA-seq expression profile is from B6 germ cells. (B) Boxplot comparing distribution of gene expression (log2 transcripts per million, tpm) with (grey, n = 606) or without (black, n = 14,419) a HELLS ChIP-seq peak at their respective promoters (mean ± SEM, *p* value – Welch's two-sided test). (C) Meta-profile of H3K4me3 (red) and HELLS (grey) signals at gencode TSS that overlap HELLS ChIP-seq peaks (n = 633). Signals are background-subtracted and scaled to maximum value to highlight positional relationship between nucleosome position and HELLS binding.

**Table S1. High throughput sequencing libraries used in this study.**

**Table S2. Oligonucleotides used for quantitative PCR in this study.**

### Figure S1

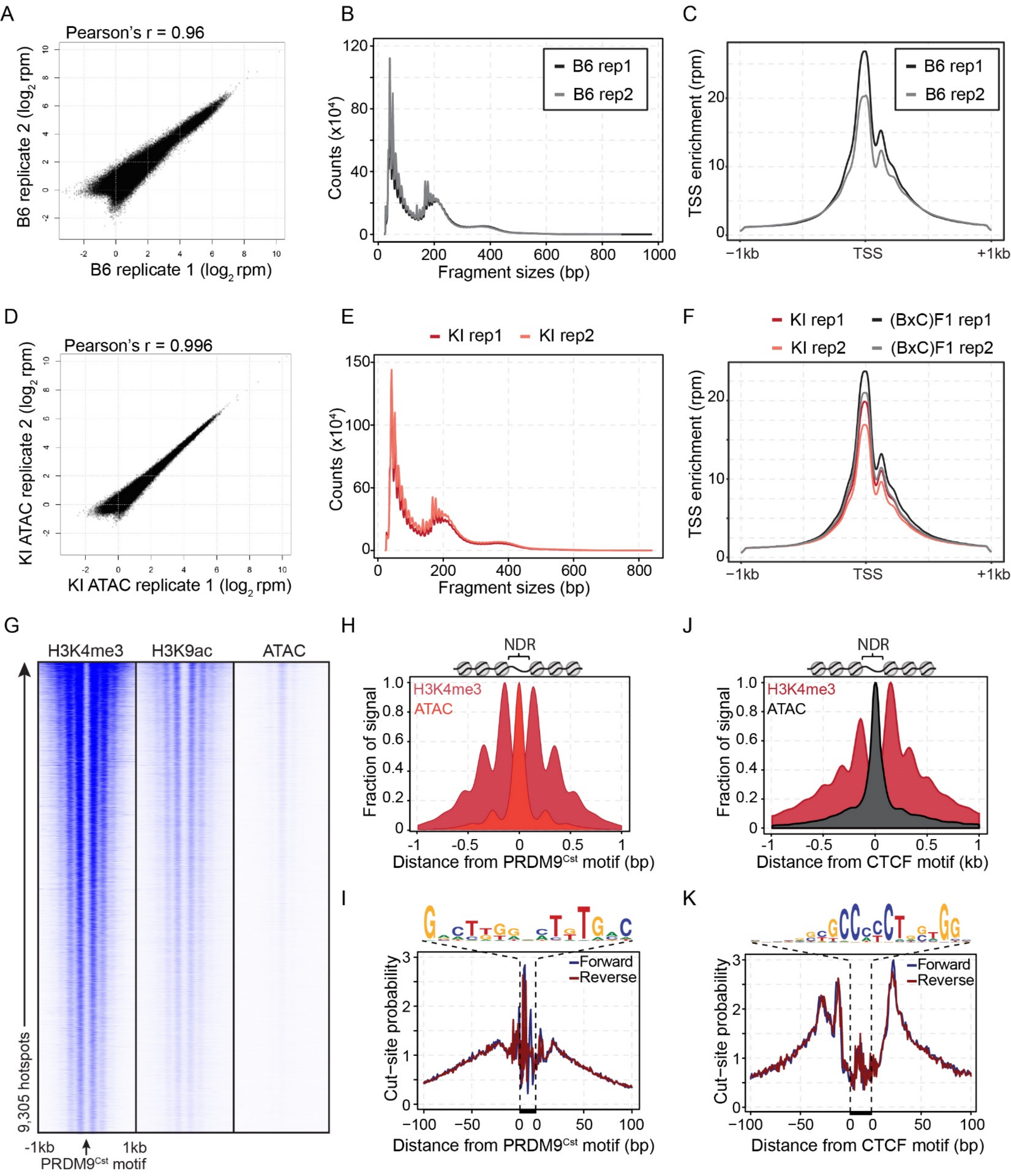

A
Pearson's r = 0.96
B6 replicate 2 (log2 rpm)
B6 replicate 1 (log2 rpm)
B
B6 rep1
B6 rep2
Counts (x10⁴)
Fragment sizes (bp)
C
B6 rep1
B6 rep2
TSS enrichment (rpm)
−1kb
TSS
+1kb
D
Pearson's r = 0.996
KI ATAC replicate 2 (log2 rpm)
KI ATAC replicate 1 (log2 rpm)
E
KI rep1
KI rep2
Counts (x10⁴)
Fragment sizes (bp)
F
KI rep1
(BxC)F1 rep1
KI rep2
(BxC)F1 rep2
TSS enrichment (rpm)
−1kb
TSS
+1kb
G
H3K4me3
H3K9ac
ATAC
9,305 hotspots
-1kb
1kb
PRDM9Cst motif
H
NDR
H3K4me3
ATAC
Fraction of signal
Distance from PRDM9Cst motif (bp)
J
NDR
H3K4me3
ATAC
Fraction of signal
Distance from CTCF motif (kb)
I
Forward
Reverse
Cut−site probability
Distance from PRDM9Cst motif (bp)
K
Forward
Reverse
Cut−site probability
Distance from CTCF motif (bp)

### Figure S2

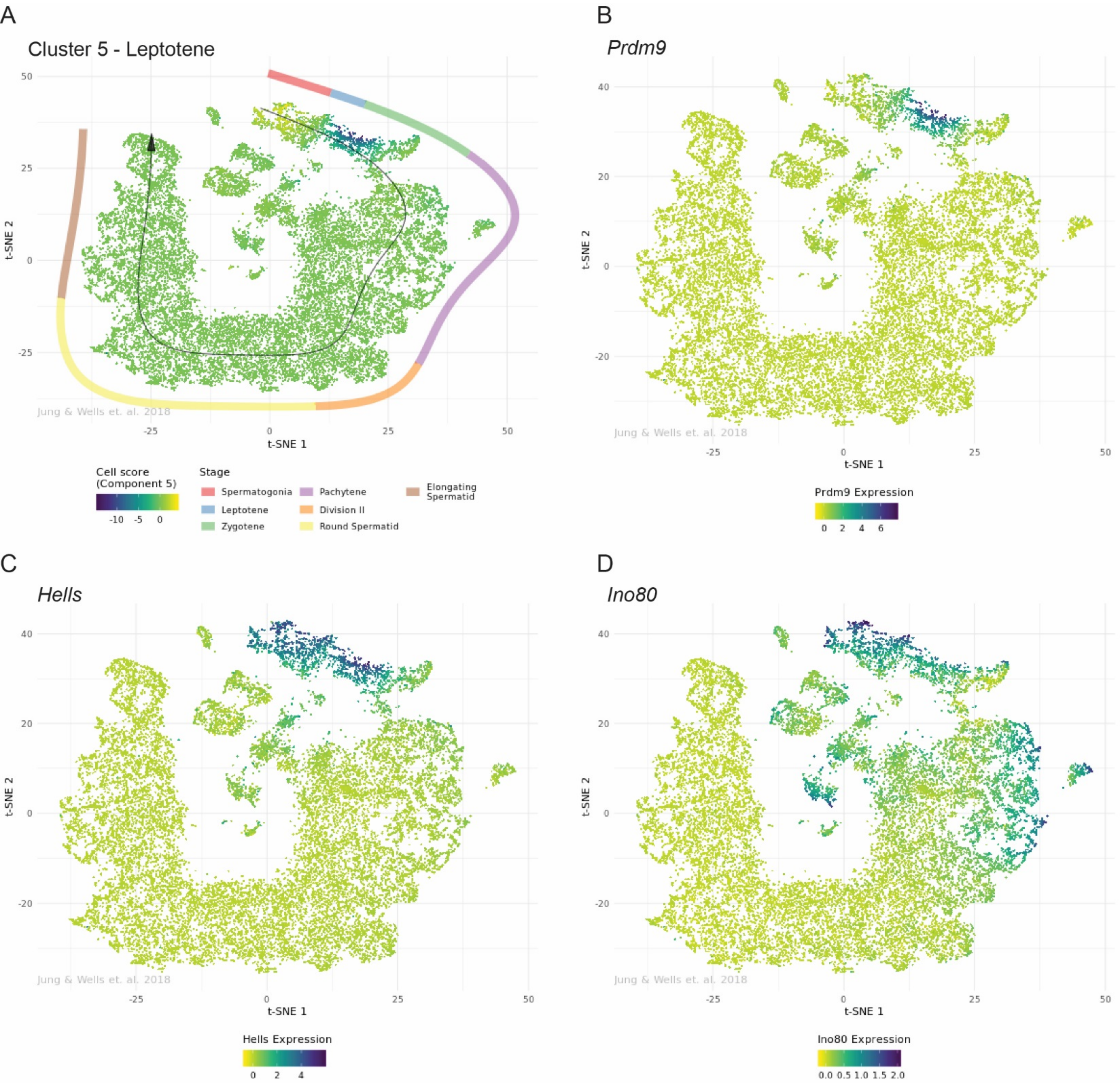
A
Cluster 5 - Leptotene
t-SNE 1
t-SNE 2
Jung & Wells et. al. 2018
Cell score (Component 5)
-10 -5 0
Stage
Spermatogonia
Leptotene
Zygotene
Pachytene
Division II
Round Spermatid
Elongating Spermatid
B
Prdm9
Prdm9 Expression
0 2 4 6
C
Hells
Hells Expression
0 2 4
D
Ino80
Ino80 Expression
0.0 0.5 1.0 1.5 2.0

Figure S3

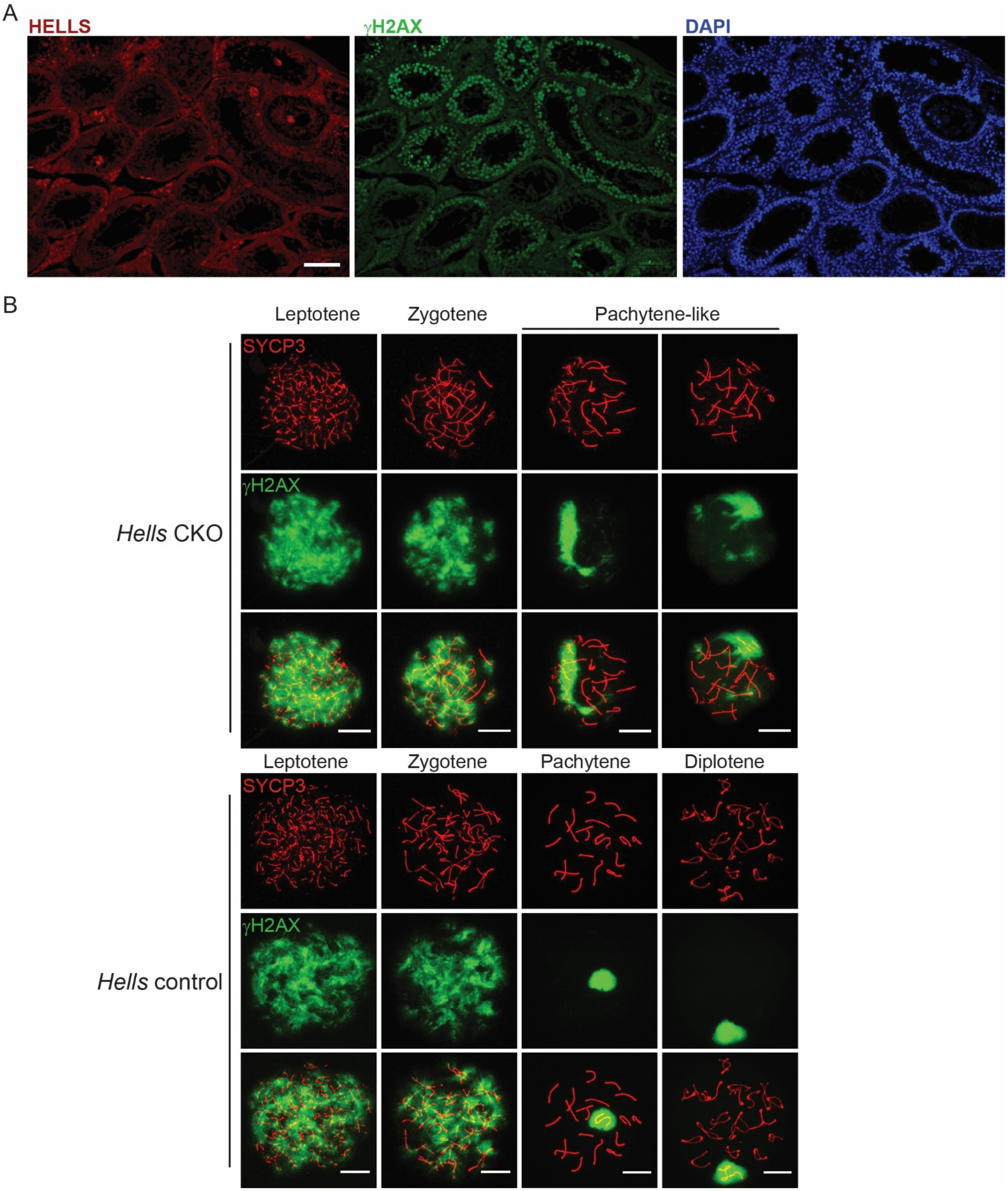
A
HELLS
γH2AX
DAPI
B
Leptotene
Zygotene
Pachytene-like
SYCP3
γH2AX
Hells CKO
Leptotene
Zygotene
Pachytene
Diplotene
SYCP3
γH2AX
Hells control

Figure S4

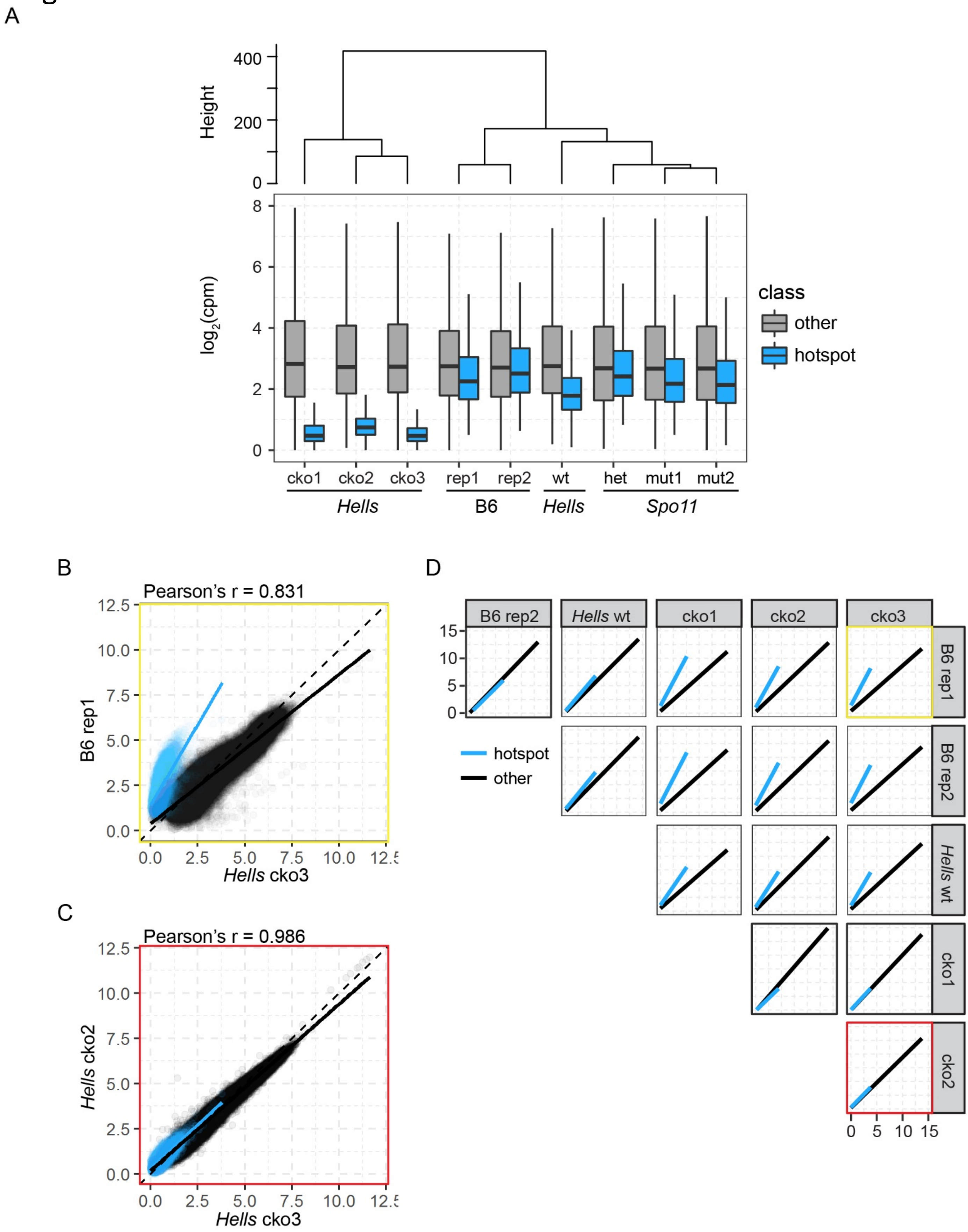
A
Height
400
200
0
log2(cpm)
8
6
4
2
0
class
other
hotspot
cko1
cko2
cko3
rep1
rep2
wt
het
mut1
mut2
Hells
B6
Hells
Spo11
B
Pearson's r = 0.831
B6 rep1
Hells cko3
C
Pearson's r = 0.986
Hells cko2
Hells cko3
D
B6 rep2
Hells wt
cko1
cko2
cko3
B6 rep1
B6 rep2
Hells wt
cko1
cko2
hotspot
other

### Figure S5

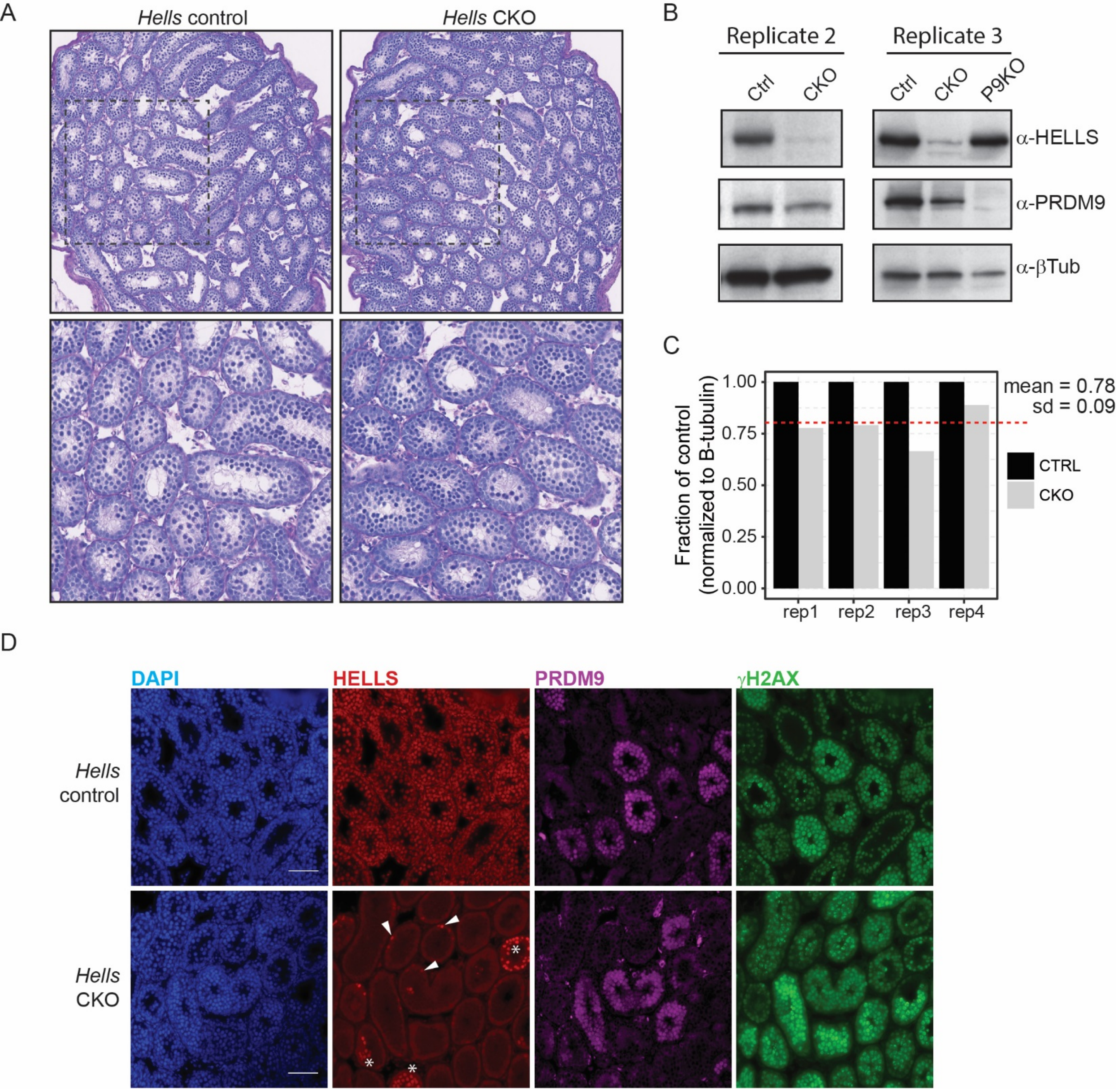
A
Hells control
Hells CKO
B
Replicate 2
Replicate 3
Ctrl
CKO
Ctrl
CKO
P9KO
α-HELLS
α-PRDM9
α-βTub
C
Fraction of control
(normalized to B-tubulin)
1.00
0.75
0.50
0.25
0.00
rep1
rep2
rep3
rep4
mean = 0.78
sd = 0.09
CTRL
CKO
D
DAPI
HELLS
PRDM9
γH2AX
Hells control
Hells CKO

Figure S6

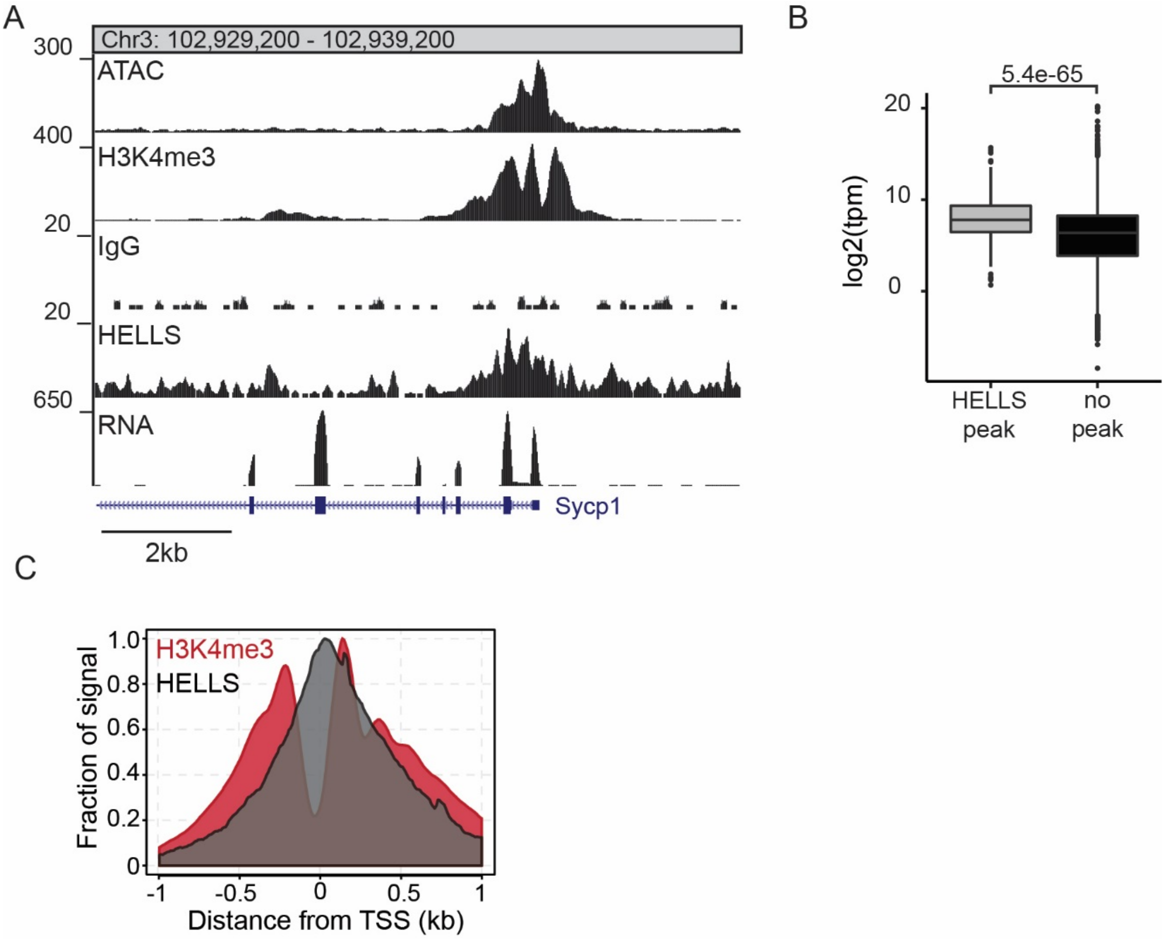

A
Chr3: 102,929,200 - 102,939,200
300
ATAC
400
H3K4me3
20
IgG
20
HELLS
650
RNA
Sycp1
2kb
B
5.4e-65
20
10
0
log2(tpm)
HELLS peak
no peak
C
H3K4me3
HELLS
1.0
0.8
0.6
0.4
0.2
0
Fraction of signal
-1
-0.5
0
0.5
1
Distance from TSS (kb)

| name | strain | antibody | age | GEO |
|---|---|---|---|---|
| ATAC4 | C57BL/6J | atac | 14 dpp | GSE135896 |
| ATAC7 | C57BL/6J | atac | 14 dpp | GSE135896 |
| ATAC13 | CAST_KI | atac | 14 dpp | GSE135896 |
| ATAC14 | CAST_KI | atac | 14 dpp | GSE135896 |
| ATAC15 | DBA/2J | atac | 14 dpp | GSE135896 |
| ATAC16 | DBA/2J | atac | 14 dpp | GSE135896 |
| ATAC17 | (B6xCAST) F1 | atac | 14 dpp | GSE135896 |
| ATAC18 | (B6xCAST) F1 | atac | 14 dpp | GSE135896 |
| ATAC56 | Spo11/+ | atac | 14 dpp | GSE135896 |
| ATAC57 | Spo11 mutant | atac | 14 dpp | GSE135896 |
| ATAC58 | Spo11 mutant | atac | 14 dpp | GSE135896 |
| ATAC61 | Hells mutant | atac | 12 dpp | GSE135896 |
| ATAC62 | Hells wt | atac | 12 dpp | GSE135896 |
| ATAC84 | Hells mutant | atac | 12 dpp | GSE135896 |
| ATAC85 | Hells wt | atac | 12 dpp | GSE135896 |
| ATAC86 | Hells mutant | atac | 12 dpp | GSE135896 |
| chip1 | Spo11 mutant | H3K4me3 | 14 dpp | GSE135896 |
| chip2 | Spo11 control | H3K4me3 | 14 dpp | GSE135896 |
| chip9 | Hells mutant | input | 12 dpp | GSE135896 |
| chip10 | Spo11 mutant | H3K4me3 | 14 dpp | GSE135896 |
| chip11 | Spo11 mutant | H3K9ac | 14 dpp | GSE135896 |
| chip15 | Hells mutant | H3K4me3 | 12 dpp | GSE135896 |
| chip16 | Hells wt | DMC1 | 6 weeks | GSE135896 |
| chip17 | Hells wt | DMC1 | 6 weeks | GSE135896 |
| chip18 | Hells wt | input | 6 weeks | GSE135896 |
| chip19 | Hells mutant | DMC1 | 6 weeks | GSE135896 |
| chip20 | Hells mutant | DMC1 | 6 weeks | GSE135896 |
| chip21 | Hells mutant | input | 6 weeks | GSE135896 |
| chip22 | Hells mutant | H3K4me3 | 12 dpp | GSE135896 |
| chip23 | Hells wt | H3K4me3 | 12 dpp | GSE135896 |
| chip34 | CAST_KI | HELLS | 12 dpp | GSE135896 |
| chip35 | CAST_KI | IgG | 12 dpp | GSE135896 |
| chip37 | CAST_KI | HELLS | 12 dpp | GSE135896 |
| chip39 | CAST_KI | input | 12 dpp | GSE135896 |
| chip40 | Hells mutant | H3K4me3 | 12 dpp | GSE135896 |
| AL2 | C57BL/6J | H3K4me1 | 12 dpp | GSE135896 |
| AM2 | C57BL/6J | H3K9me2 | 12 dpp | GSE61613 |
| AN2 | C57BL/6J | H3K4me1 | 12 dpp | GSE135896 |
| AO | C57BL/6J | H3K9me2 | 12 dpp | GSE61613 |
| AX2 | C57BL/6J | H3K27me3 | 14 dpp | GSE135896 |
| BR | (B6xCAST) F1 | H3K9ac | 12 dpp | GSE135896 |
| BV2 | C57BL/6J_NaBut | H3K9ac | 14 dpp | GSE135896 |
| BW2 | C57BL/6J_NaBut | H3K27ac | 14 dpp | GSE135896 |
| BY | C57BL/6J_NaBut | H2B120ac | 14 dpp | GSE135896 |
| CB2 | C57BL/6J | H3K27me3 | 14 dpp | GSE135896 |
| CF | C57BL/6J | H2A.Z | 14 dpp | GSE135896 |

| | | | | |
|---|---|---|---|---|
| CG2 | C57BL/6J | H3K9ac | 14 dpp | GSE135896 |
| CH2 | C57BL/6J_NaBut | H3K9ac | 14 dpp | GSE135896 |
| CI2 | C57BL/6J_NaBut | H3K27ac | 14 dpp | GSE135896 |
| CJ | CAST_KI | H3K9ac | 14 dpp | GSE135896 |
| CK | CAST_KI | H3K9ac | 14 dpp | GSE135896 |
| hui1 | C57BL/6J | H3K9me3 | 14 dpp | GSE61613 |
| natalie1 | C57BL/6J | H3K36me3 | 14 dpp | GSE76416 |
| germ_B6-rep3 | C57BL/6J | H3K4me3 | 14 dpp | GSE113192 |
| germ_D2-rep1 | DBA/2J | H3K4me3 | 14 dpp | GSE113192 |
| B6xKI_H3K4me3_ChIP_sample_[illegible] | (B6xCAST) F1 | H3K4me3 | 12 dpp | GSE52628 |
| B6xKI_H3K4me3_ChIP_sample_[illegible] | (B6xCAST) F1 | H3K4me3 | 12 dpp | GSE52628 |
| KI_H3K4me3_ChIP_sample_1 | CAST_KI | H3K4me3 | 12 dpp | GSE52628 |
| KI_H3K4me3_ChIP_sample_2 | CAST_KI | H3K4me3 | 12 dpp | GSE52628 |

| Purpose | Name | Sequence |
|---|---|---|
| Hells genotyping | Hells_1 | GTAAGAGTCTCAGTGTCAACC |
| | Hells_2 | CAACGGGTTCTTCTGTTAGTCC |
| | Hells_3 | AAGTCGTCGTCCTTACCAGTG |
| | Hells_4 | AGGACTCCAGGCAAATCTGA |
| ChIP-qPCR | C1_114_peak_1F | CTGCCTTCATTCCACTCCTC |
| | C1_114_peak_1R | CAGGGAGGGAAAACATAAATCA |
| | C1_114_valley_2F | TGTTCCACACCCAGCTATTG |
| | C1_114_valley_2R | ATCATGGGGCAAGATCAAAC |
| | C2_44_peak_1F | CTTGTAGAACTGAGATTAGTTGAGAGC |
| | C2_44_peak_1R | GCTTTTCCTGTTTCTCCCCTA |
| | C2_44_valley_2F | CACTGGGGATGGTAGCATTAG |
| | C2_44_valley_2R | TCAAGGACACAGGGGATGAT |
| | C3_F | ATTGCTAGAAAGGCGTGTGC |
| | C3_R | TTGCTAGGCATGTGAAATGG |
| | A1_1F | AACTGCACAGCTGCAAACAC |
| | A1_1R | TATCCCAACCAATCCCATGT |
| | hGAPDH_F | GAGCCTCGAGGAGAAGTTCC |
| | hGAPDH_R | GACTGAGATGGGGAATTGGA |

| notes |
| --- |
| to genotype tm1a use Hells_1 and Hells_2; expected size 663bp |
| to genotype tm1c use Hells_3 and Hells_4; wt expected size 323 bp mutar |
| to genotype tm1d use Hells_1 and Hells_4; expected mut size 705 bp |
| Figure 6B, 6C |
| Figure 6B, 6C |
| Figure 6C, 6H, 7A |
| Figure 6C, 6H, 7A |
| Figure 6B, 6C |
| Figure 6B, 6C |
| Figure 6C, 6H, 7A |
| Figure 6C, 6H, 7A |
| Figure 6H, 7A |
| Figure 6H, 7A |
| Figure 6C, 6H, 7A |
| Figure 6C, 6H, 7A |
| Figure 6C, 6H, 7A |
| Figure 6C, 6H, 7A |

ıt size 385 bp
